# Deciphering tumor ecosystems at super-resolution from spatial transcriptomics with TESLA

**DOI:** 10.1101/2022.11.05.515256

**Authors:** Jian Hu, Kyle Coleman, Edward B. Lee, Humam Kadara, Linghua Wang, Mingyao Li

**Affiliations:** Department of Human Genetics, School of Medicine, Emory University, Atlanta, GA 30322, USA; Department of Biostatistics, Epidemiology and Informatics, Perelman School of Medicine, University of Pennsylvania, Philadelphia, PA 19104, USA; Translational Neuropathology Research Laboratory, Department of Pathology and Laboratory Medicine, Perelman School of Medicine, University of Pennsylvania, Philadelphia, PA 19104, USA; Department of Translational Molecular Pathology, The University of Texas MD Anderson Cancer Center, Houston, TX 77030, USA; Department of Genomic Medicine, The University of Texas MD Anderson Cancer Center, Houston, TX 77030, USA; The University of Texas MD Anderson Cancer Center UTHealth Graduate School of Biomedical Sciences (GSBS), Houston, TX 77030, USA

**Keywords:** Spatial transcriptomics (ST), Tumor-infiltrating lymphocytes (TILs), Tumor microenvironment (TME), Tertiary lymphoid structure (TLS)

## Abstract

Recent advances in spatial transcriptomics (ST) have enabled the comprehensive characterization of gene expression in tumor microenvironment. However, ST only measures expression in discrete spots, which limits their usefulness in studying the detailed structure of TME. Here we present TESLA, a machine learning framework for multi-level tissue annotation in ST. TESLA integrates histological information with gene expression to annotate heterogeneous immune and tumor cells directly on the histology image, and further detects tertiary lymphoid structures and differential transcriptome programs between the edge and core of a tumor. TESLA provides a powerful tool for understanding the spatial architecture of the TME.

## Introduction

The tumor microenvironment (TME) contains networks of cells and structures that surround tumor cells [1]. Tumor-TME crosstalk regulates the initiation, progression, and metastasis of tumors [2]. A comprehensive analysis of the multiple exchanges between tumor cells and their TME is essential for understanding the underlying mechanisms of tumor growth and response to therapy. The TME has a great impact on the efficacy of anti-cancer therapies and clinical outcomes [3, 4]. For example, hypoxic avascular regions deeply embedded inside the tumors significantly hinder the delivery of therapeutic agents[5], and hypoxic tolerant tumor cells are resistant to most anti-cancer treatments [6, 7].

Tumor infiltrating lymphocytes (TILs) consist of lymphocytic cell populations that have invaded the tumor tissue and have emerged as an important biomarker in predicting the efficacy and outcome of anti-cancer treatment [8, 9]. Recent studies suggest that not only the density of TILs, but also their spatial organization, such as the presence of tertiary lymphoid structures (TLSs), play key roles in determining tumor immune phenotypes and response to immunotherapy [10]. TLSs are broadly found in the TME for various solid cancers. Similar to secondary lymphoid organs, mature TLSs are believed to be the site of immune response activation against tumor by recruiting and activating TILs and represent the key to understanding antitumor immune responses.

Spatial transcriptomics (ST) enables gene expression profiling while preserving location information in tissues, which innovates a promising avenue to understand the spatial context and the nature of cellular heterogeneity of the TME [11-13]. In a ST experiment using spatial barcoding followed by next-generation sequencing-based technologies such as Spatial Transcriptomics [14] and 10x Genomics Visium, the expression levels for thousands of genes in a tissue section are simultaneously measured, complemented by a high-resolution histology image obtained from the same tissue section. With the power of ST, we aim to provide a detailed annotation of tumor structure and different lymphocytes by integrating gene expression and histology image information.

A major challenge that hinders gene expression and histology integration in spatial barcoding-based ST is the relatively low resolution of gene expression data compared to histology images, which prevents deciphering detailed TME structures such as the TLS. Although methods such as BayesSpace [15] can enhance the gene expression resolution, their enhancement is only for regions captured by spots, but still leaves a large portion of the tissue unmeasured for gene expression. A recently developed method, XFuse [16], uses a deep generative model to predict super-resolution gene expression at each pixel by leveraging information in hematoxylin and eosin (H&E)-stained histology images. However, XFuse is extremely slow – often takes more than two weeks to analyze a tissue section generated from 10x Visium on a GPU machine. Additionally, its prediction accuracy heavily depends on the similarity between the spatial patterns of gene expression and the histology images. For many genes that are involved in innate and adaptive immune responses, their spatial patterns are weakly correlated with histology images, making XFuse less applicable when the goal is to decipher the TME.

To overcome the above-mentioned challenge, we present TESLA (<u>T</u>umor <u>E</u>dge <u>S</u>tructure and <u>L</u>ymphocyte multi-level <u>A</u>nnotation), a machine learning framework that integrates gene expression and histology image information in ST to investigate the TME. By generating super-resolution gene expression images, TESLA can leverage histology information to annotate different tumor/TME cell types on the histology image with pixel-level resolution, detect TIL structure such as TLS by colocalization analysis of different lymphocytes and dendritic cells, and characterize high-resolution cellular and molecular spatial structures of tumor by separating tumor edge and core and elucidating differential transcriptome programs between the edge and core. The detailed multi-level annotations performed by TESLA provide a comprehensive understanding of the TME.

## Results

### Overview of TESLA and evaluation

TESLA starts from enhancing the original spot-level gene expression in ST. This step is based on two assumptions: 1) spots that are physically close have similar gene expression, and 2) the expression patterns for spatially variable genes are correlated with histology image features. As shown in **Fig. 1a**, TESLA first detects the contour of the tissue area. Next, for each gene, TESLA imputes each superpixel’s gene expression through weighted aggregation of spot-level expression values from nearby spots and generates an expression image for the entire histology image by filling in tissue gaps that are not covered by spots.

**Fig. 1.**
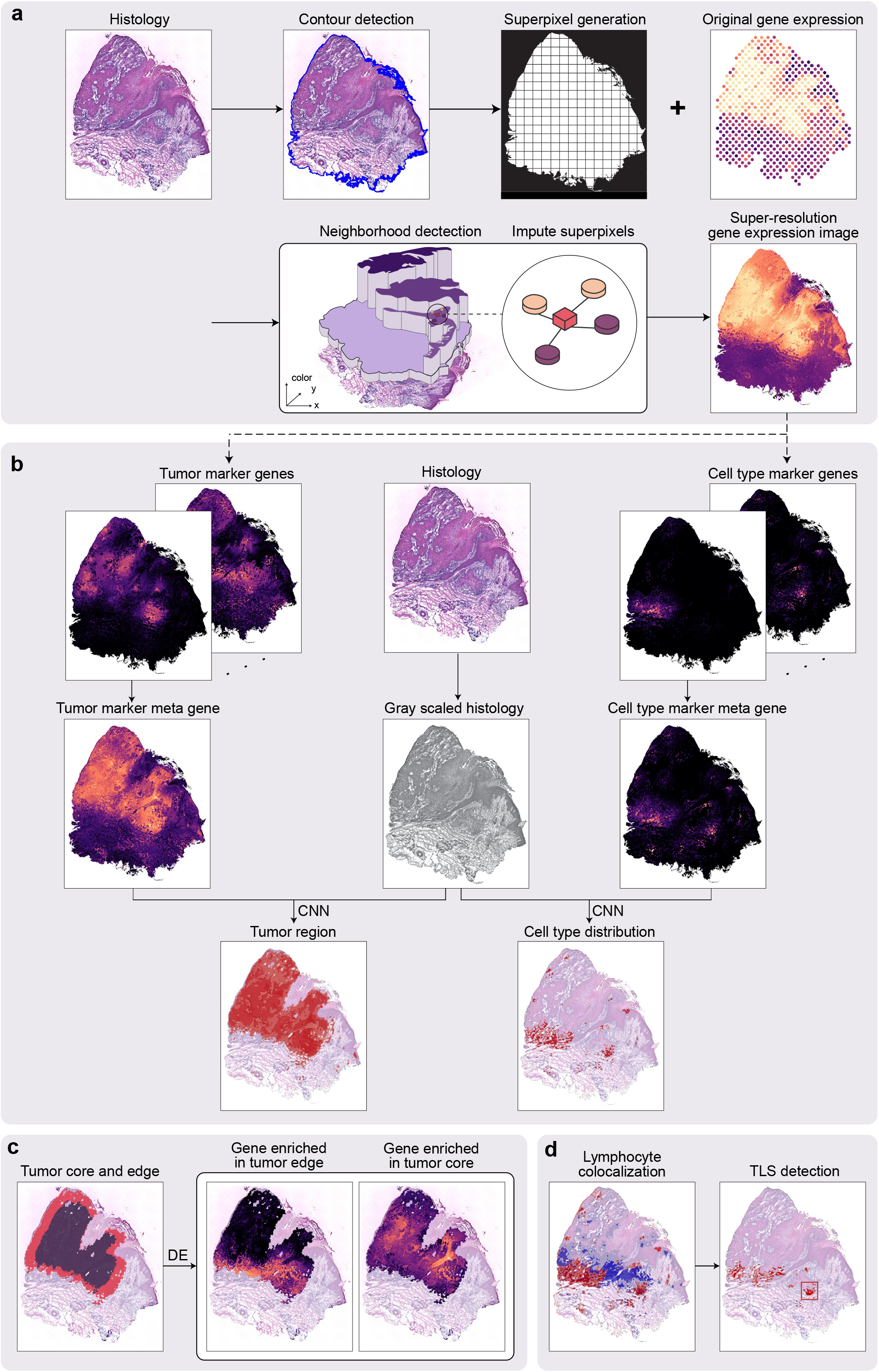
Workflow of TESLA. **a**, TESLA starts from generating superpixels to cover the entire tissue area. The gene expression of each superpixel is imputed by the weighted sum of its nearest measured spots in a 3-D space, defined by the spatial coordinates (*x, y*) and the 3-rd dimensional coordinate *z*, obtained from the RGB values in the histology image. **b**, TESLA combines a set of tumor/cell type marker genes into a meta gene and stacks the meta gene image with the gray-scaled histology image into a two-channel image, which is fed into a convolutional neural network (details shown in **Fig. 6**) for segmentation. Tumor region/cell type distribution can be displayed on the histology image based on the segmentation results. **c**, TESLA can further separate the tumor region into edge and core. Downstream differential expression analysis is performed to identify core or edge enriched genes. **d**, Based on the colocalization of specific lymphoid cell types detected in **b**, TESLA can identify TLS on the histology image.

With the super-resolution gene expression images generated for tumor, immune, and stromal cells, TESLA can annotate the tumor region and tumor-immune interface at multiple levels. To infer tumor cell distribution, TESLA first combines tumor related marker genes into a meta gene that captures the shared expression patterns of the markers **(Fig. 1b)**. After stacking the meta gene images and the histology image as input, TESLA then employs a convolutional neural network approach to segment the tissue region in an unsupervised manner, which enables the annotation of the tumor region directly on the histology image with pixel-level resolution. Since the tumor cells and microenvironment at the invasive edge of tumor are known to be different from that of the tumor core [2], TESLA also draws the leading edge along the tumor boundary and outlines the tumor edge and tumor core **(Fig. 1c)**.

To characterize spatial cellular and molecular heterogeneity between the tumor edge and core and better understand tumor-TME dynamics, TESLA first performs differential gene expression analysis between the edge and core, which enables the identification of differentially expressed genes (DEGs) that are specific to the tumor edge or core regions. With the region-specific DEGs, TESLA can also compare cell compositions and transcriptome programs between the edge and core and further profile spatial heterogeneity of the TME. In addition to tumor region annotation, TESLA can also annotate a target cell type’s distribution on the histology image by using cell-type-specific marker genes, e.g., lymphoid cell lineage markers, as input **(Fig. 1b)**. Based on the colocalization of specific cell populations, TESLA can further detect multicellular structures such as TLS **(Fig. 1d)**.

To showcase the strength and properties of TESLA, we applied it to eight publicly available ST datasets [15, 17-23] **(Supplementary Table 1)**. First, we show that the super-resolution gene expression correlates well with protein expression obtained from immunofluorescence staining. We further show that the cell type distribution predicted by TESLA agrees well with independent cell type deconvolution results by RCTD[24], and the predicted TLSs are in agreement with independent pathologist’s annotation.

### Imputation of super-resolution gene expression by integrating histology image information

TESLA’s multi-level cellular and molecular annotation of ST data relies on the super-resolution gene expression images. We conducted hold-off experiments on five public datasets [15, 17, 18, 22, 23] to show that TESLA can faithfully recover the missing expression patterns for genes with spatial variability. For each dataset, we first selected Spatially Variable Genes (SVGs) using our previously developed method, SpaGCN [25]. Next, we masked each measured spot and used the remaining spots to impute the SVGs’ expression for the masked spot. We repeated this masking for all of the spots and examined whether TESLA can faithfully impute the missing gene expression at the masked spots. **Fig. 2a** shows the correlation between the observed and imputed SVGs’ expression of the masked spots. The median correlation among the SVGs is 0.60 for the invasive ductal carcinoma (IDC) dataset [15], 0.71 for the cutaneous squamous cell carcinoma (CSCC) dataset [17], 0.69 for the cutaneous malignant melanoma dataset [18], 0.70 for the mouse posterior brain tissue dataset, and 0.54 for the mouse kidney dataset. These high correlations indicate that TESLA can recover the underlying expression pattern for unmeasured spots. Since the distances between superpixels and their nearest neighboring spots in real ST data are smaller than the distances between the masked spots and their neighboring spots in the hold-off experiments, we expect TESLA to perform better in real data analysis. We also performed mask test in the same way using the top 2,000 highly variable genes and the results are shown in **Supplementary Fig. 1**.

**Fig. 2.**
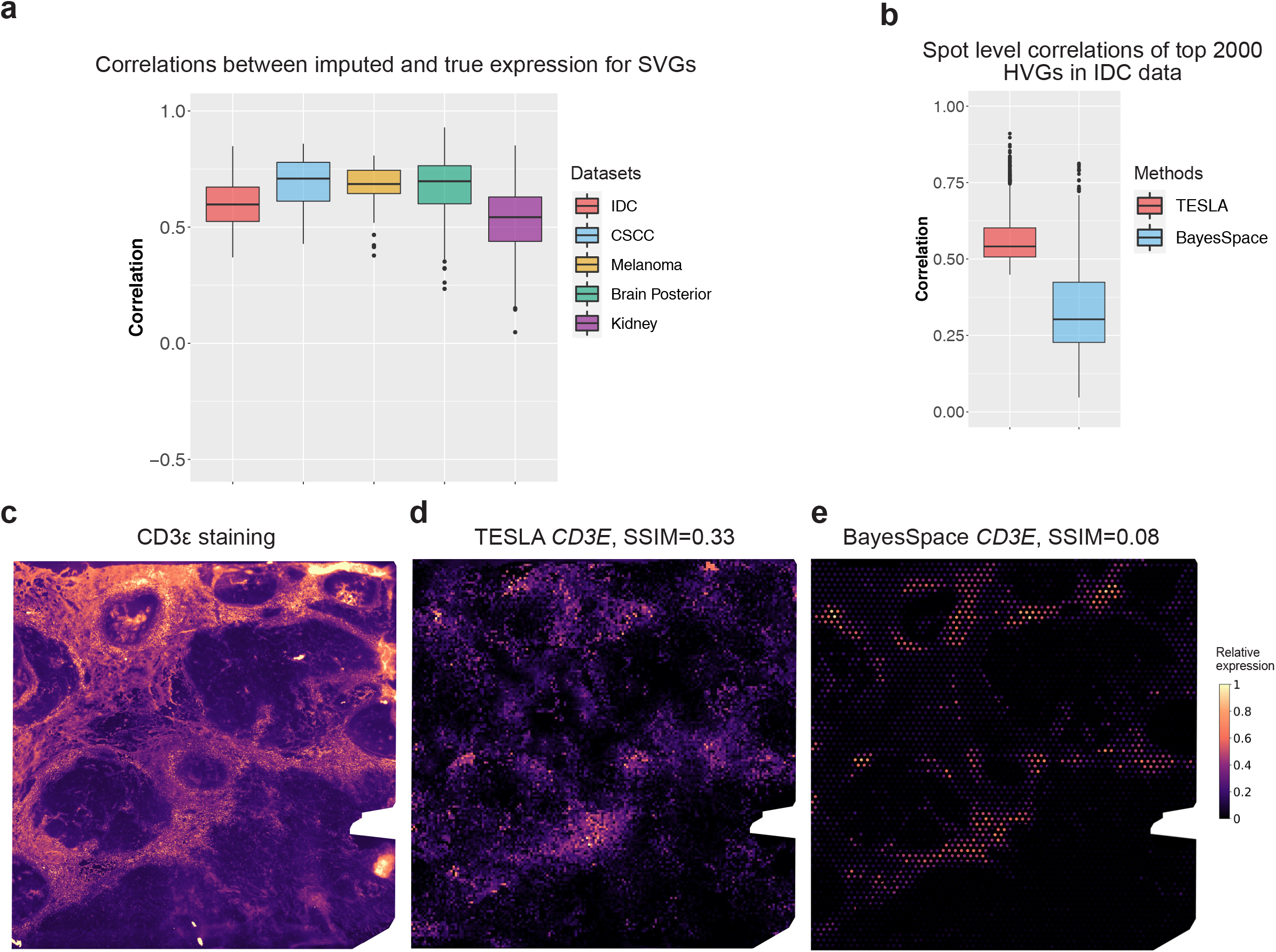
Evaluation of super-resolution gene images from TESLA. **a**, Boxplot of Pearson correlations between the original gene expression and TESLA’s imputed gene expression for masked spots for SVGs detected by SpaGCN. Each point in the boxplot represents a gene. SpaGCN detected 68 SVGs (n=68) from the IDC dataset, 85 SVGs (n=85) from the CSCC dataset and 30 SVGs (n=30) SVGs from the cutaneous malignant melanoma dataset. The lower and upper hinges correspond to the first and third quartiles, and the center refers to the median value. The upper (lower) whiskers extend from the hinge to the largest (smallest) value no further (at most) than 1.5 × interquartile range from the hinge. Data beyond the end of the whiskers are plotted individually. **b**, Boxplot of Pearson correlations between the original spot-level gene expression and “spot-level” gene expression obtained from the enhanced expression generated from TESLA (n=2,000) and BayesSpace (n=2,000) for the top 2,000 highly variable genes selected by BayesSpace from the IDC dataset. Each point in the boxplot represents a gene. Boxplot hinges, median, and whiskers are defined the same was as in **a. c**, Intensity of the CD3ε immunofluorescence staining. Intensity was scaled to (0, 1) for visualization. **d**, Super-resolution gene expression image of *CD3E* from TESLA. **e**, Super-resolution gene expression image of *CD3E* from BayesSpace.

To further evaluate the accuracy of TESLA’s imputed super-resolution gene expression, we performed additional analysis on the IDC dataset, which is generated from an estrogen receptor-positive (ER+), progesterone receptor-negative (PR-), human epidermal growth factor receptor (HER)2-amplified (HER+) invasive ductal carcinoma tissue section [26]. This tissue section has paired immunofluorescence staining for CD3ε protein counterstained with 4,6-diamidino-2-phenylindolo (DAPI). We compared the performance of TESLA with BayesSpace [15], a recently developed tool that can also enhance the gene expression resolution for ST data. TESLA features two benefits relative to BayesSpace because 1) TESLA can incorporate high-resolution histology image information and offers more flexibility for resolution enhancement by adjusting the superpixel size, while BayesSpace can only enhance the gene expression resolution by splitting a spot into fixed number of subspots; 2) TESLA can fill in gene expression for unmeasured tissue regions between spots while BayesSpace leaves unmeasured tissue areas blank. For 10x Visium, each spot is 55 μm in diameter with a 100 μm center to center distance between spots. The lack of full tissue coverage by spots leaves approximately 54% to 80% of the tissue unmeasured for gene expression **(Supplementary Note 1)**.

An ideal gene expression enhancement method should increase the gene expression resolution while retaining the original expression pattern at the spot level as this will ensure no artificial patterns are introduced in the enhanced gene expression. To show that TESLA outperforms BayesSpace in retaining the original gene expression patterns, we considered the top 2,000 highly variable genes selected by BayesSpace and obtained the spot-level gene expression from the enhanced expression generated from each method. For TESLA, we obtained the spot-level gene expression from the super-resolution gene expression image by extracting expression from a circle that exactly overlaps the measured spots. For BayseSpace, we summed up the expression from all sub-spots within a spot to get the spot-level expression. As shown in **Fig. 2b**, TESLA’s super-resolution gene expression derived spot-level expression yields significantly higher correlations with the original spot-level gene expression than BayesSpace (median: 0.54 vs 0.30, two-sample t-test *P* < 2.2e-16). Although BayesSpace requires the sum of principal components of the subspots equals the principal component of the original spot, it cannot guarantee the expression pattern is not distorted in the original gene expression space **(Supplementary Fig. 2)**.

Since the ground truth of gene expression is unknown, we cannot evaluate the accuracy of the super-resolution gene expression directly. Instead, we evaluated the super-resolution gene expression with the immunofluorescence staining for the same gene-protein pair obtained from the same tissue section in the IDC dataset, which includes immunofluorescence staining for CD3ε, a T cell co-receptor that is involved in activating both the cytotoxic T cell and T helper cells. An accurate super-resolution gene expression imputation approach should yield gene expression that correlates with the protein expression obtained from immunofluorescence staining [15]. We treated the protein expression of CD3ε staining as the ground truth **(Fig. 2c)** and evaluated the enhanced gene expression at the pixel level by comparing their similarity with the immunofluorescence image. We utilized Structural Similarity Index (SSIM) [27], a commonly used metric for measuring the similarity between two images, for evaluation. SSIM ranges from -1 to 1 with a value of 1 indicating two identical images, 0 indicating no structural similarity, and -1 indicating strong dissimilarity. As shown in **Fig. 2d**, the SSIM between the protein staining image and the super-resolution *CD3E* gene image from TESLA is 0.33, while the SSIM is only 0.08 for BayesSpace (**Fig. 2e**), presumably due to its inability to fill in gene expression in the unmeasured tissue area. We also compared TESLA with BayesSpace using root mean squared error and reached a similar conclusion **(Supplementary Fig. 3)**.

### Characterization of high-resolution cellular and molecular spatial structure of tumor

The tumor edge is considered as the tumor invasion front due to its strong invasive ability and it has been reported that tumor edge has unique microenvironment and distinct morphological, structural, and molecular features from that of tumor core, such as the level of immune cell infiltration and compositions, vascular density, hypoxia status, metabolic rate, etc. [28, 29]. The super-resolution gene expression images generated by TESLA enables the detection and precise annotation of tumor edge, which enables further exploration of the structural, cellular, and molecular differences in tumor and TME cells between the primary tumor core and edge.

To show that such analyses can advance our understanding of intra-tumor heterogeneity, we first analyzed the CSCC dataset of the skin [17]. As reported in the original study, the tumors include basal, cycling, and differentiating keratinocyte cell populations that are similar to normal skin, and a tumor-specific keratinocyte (TSK) population. Therefore, we identified the tumor region using nine TSK marker genes suggested in the original study (*NT5E, PTHLH, INHBA, LAMC2, MMP10, FEZ1, CD151, IL24, SLITRK6*), plus three commonly used marker genes (*TOP2A, KIF1C, BUB1B*). Utilizing these genes **(Supplementary Fig. 4)**, TESLA derived a meta gene that captured all their patterns and further detected the tumor region by joint segmentation of the meta gene and histology images. **Fig. 3a** shows that the tumor region detected by TESLA agrees well with the dark region in the histology image except the upper right and bottom right regions. Further examination of the histology image of the upper right region by a pathologist (E.B.L.) revealed that this region consists of dead cells with hypereosinophilic cytoplasm and pkynotic nuclei (yellow box). This is further confirmed by the relatively low total unique molecular identifier (UMI) counts compared to other regions **(Supplementary Fig. 5)**. The region on the right represents non-neoplastic skin adjacent to the squamous cell carcinoma (blue box). This example shows that by combining gene expression and histology image together, TESLA can accurately annotate live tumor region from non-viable or non-neoplastic tissue without the reliance on a pathologist’s manual annotation.

**Fig. 3.**
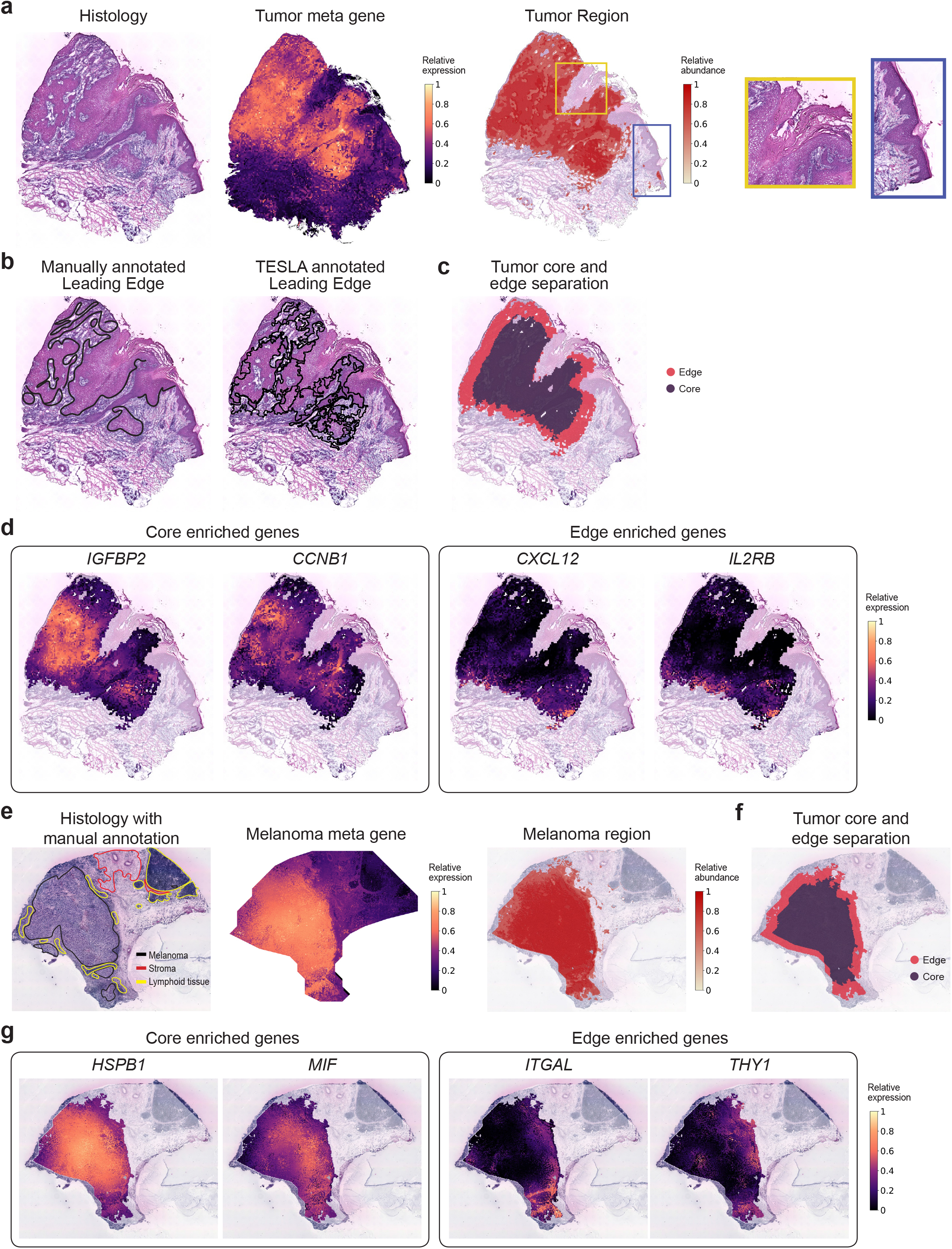
TESLA characterizes intra-tumor heterogeneity in a CSCC sample and a melanoma sample. **a**, Histology image (left), tumor meta marker gene (middle) and the tumor region detected by TESLA (right) from the CSCC tissue section. The yellow and blue boxes mark two regions with dark color on histology image but not detected as tumor region. **b**, Pathologist’s manually annotated leading edge from the original study and the leading edge drawn by TESLA for the CSCC tissue section. **c**, Tumor core and edge separation obtained by TESLA for the CSCC tissue section. **d**, Examples of genes enriched in the tumor core (*IGFBP2, CCNB1*) and tumor edge (*CXCL12, IL2RB*) detected by TESLA for the CSCC tissue section. **e**, Pathologist annotated histology image from original study, melanoma meta marker gene expression, and the melanoma region detected by TESLA. **f**, tumor core and edge separation of the melanoma region by TESLA. **g**, Examples of the tumor core (*HSPB1, MIF*) and edge (*ITGAL, THY1*) enriched genes detected by TESLA for the melanoma tissue section.

The leading edge drawn by TESLA also agrees well with pathologist’s manual **annotation (Fig. 3b)**. After detecting the tumor boundary, TESLA further separated the tumor into core and edge **(Fig. 3c)**. The tumor edge is defined as the area that is within 200 μm interface of tumor and adjacent non-malignant tissue and the tumor core is defined as the proximal tumor area within the tumor edge[28]. DEG analysis between the tumor core and edge identified genes that are highly enriched in the core (3,665 genes) or edge (106 genes). For example, in **Fig. 3d** and **Supplementary Fig. 6**, oncogenes such as *IGFBP2, CCNB1, KRT15* and *HRAS* are highly expressed in the core region, whereas tumor edge enriched genes included *CXCL12, IL2RB, LAIR1*, and *LILRB2*, which are immune-related, implying a distinct microenvironment from that of the tumor core. We further performed gene set enrichment analysis for the 106 tumor edge enriched genes and the top 300 tumor core enriched genes with the highest fold changes, by computing overlaps with curated gene sets in the Molecular Signature Database [30] **(Supplementary Table 2)**. Our analysis suggested significant enrichment of the metabolism of lipids under hypoxic conditions and mitotic phase in cell cycle in the tumor core. The high proliferation rate of cancer cells in the tumor core leads to hypoxia and nutrient deprivation, which affect the lipid metabolism of cells [31, 32]. Pathways that are highly activated in the tumor edge included innate immune system, adaptive immune system, interferon-γ response, inflammatory response, and cytokine signaling. These highly activated immune systems in the tumor edge are possibly due to a higher level of immune cell density and tumor-immune interactions in the edge. Together, TESLA’s unique feature to characterize and profile tumor edge and core may provide great insights into better understanding of the TME and the mechanisms of immune suppression and evasion.

To show the generalizability of TESLA in characterizing the cellular and molecular spatial structure of tumor, we next analyzed the cutaneous malignant melanoma dataset [18]. Using a list of genes for clinical melanoma diagnosis [33] (*MITF, CSPG4, MAGEA1, MLANA, TYR* and *SOX10*; **Supplementary Fig. 7**), TESLA identified the melanoma region, which strongly recapitulate the pathologist’s manual annotation in **Fig. 3e**. Following similar steps as described previously, we further separated the tumor region into tumor core and edge **(Fig. 3f)** and detected 3,510 genes enriched in the tumor core and 155 genes enriched in the tumor edge. Some hypoxia related genes [34, 35] (e.g., *HSPB1, MIF, TPI1*) were detected to be highly expressed in the tumor core while most genes highly expressed in the tumor edge are immune related (**Fig. 3g** and **Supplementary Fig. 8**). Next, we performed gene set enrichment analysis for the 155 edge enriched genes and the top 300 core enriched genes with the highest fold changes **(Supplementary Table 3)**. Interestingly, for the top 300 core enriched genes, we observed a significant upregulation of the signaling by Rho GTPases, receptor tyrosine kinase, and Wnt, and MET activates RAS signaling, which are known to be involved in melanoma progression through regulating cell proliferation and invasion [36, 37]. In addition, we observed increased signaling by MET, which has emerged as a paradigm of tumor resistance to modern targeted therapies, and the assessment of its expression in patients’ samples may be a valuable biomarker of tumor progression and response to targeted therapy [38]. The enriched genes in the tumor edge are largely related to inflammatory response, such as *PDCD1, IL10RA, PAX5, CCL19, TNFSF4, IFI27, IL32, and IL4R*. Consistently, pathway enrichment analysis revealed increased expression of interferon gamma response, cytokine signaling and cytokine-cytokine receptor interaction, and increased signaling by Interleukins, suggesting activated anti-tumor immune response. Interestingly, we also observed up-regulated expression of genes defining epithelial-mesenchymal transition (EMT). EMT is a process through which epithelial tumor cells acquire mesenchymal phenotypic properties, and it contributes to both metastatic dissemination and therapy resistance in cancer [39]. The cell type distributions inferred by cell type deconvolution analysis with RCTD also showed significant difference between the tumor edge and core **(Supplementary Fig. 9)**, with the tumor edge enriched for T cells whereas the core enriched for malignant cells.

We also demonstrate that all these findings, based on the super-resolution annotation, cannot be achieved with the original spot-level data **(Supplementary Note 2)**. The failure of detecting tumor edge enriched genes at the spot level is possibly due to two reasons. First, the number of observations is much smaller when considering a spot as the analysis unit, and the reduced sample size in DEG analysis will lead to less power. Second, the spot-level data do not have single-cell resolution. Indeed, the diameter size of each spot is 100 μm in Spatial Transcriptomics, which is much larger than a single cell. Since each spot may contain many cells, the mixture of cells from different cell types will dilute the differential expression signal, especially when the immune cells are rare in the tumor edge. Numerous studies have shown that the tumor-immune interface plays an important role in understanding the TME with primary relevance to the success of immunotherapy [40]. Therefore, we believe that until the sequencing-based spatial transcriptomics technologies reach to single-cell resolution, gene expression resolution enhancement will be needed when the goal is to detect gene expression changes that occur only in a small region of the tissue.

### Identification of lymphoid cell types and TLS

Next, we show that TESLA can profile the spatial distribution of different immune cell subsets based on super-resolution expression images of lineage-specific genes. We first analyzed the CSCC data [17]. Using canonical cell type marker genes **(Supplementary Fig. 10)**, we detected the locations for B cells, CD4+ T cells, dendritic cells, and CXCL13, a marker of TLS that is known to be also constitutively expressed in secondary lymphoid tissue. As shown in **Fig. 4a**, TESLA’s cell type distribution has much higher resolution than results obtained from spot-level gene expression by SpaGCN [25], a clustering method developed for spatial transcriptomics data **(Fig. 4b)**. For Spatial Transcriptomics and Visium data, each spot may contain multiple cells. By leveraging high-resolution histology image data, TESLA’s cell type annotation achieves pixel-level resolution, which better describes the underlying cell type distributions. By colocalizing B cells, CD4+ T cells, dendritic cells, and CLCX13 near the tumor region, TESLA further detected TLSs **(Fig. 4c)**. Examination of the histology image by a pathologist (E.B.L.) indicates that the blue boxed region contains aggregates of lymphocytes consistent with a TLS on the edge of the tumor. We also observed the colocalization of lymphocytes in the yellow boxed region that are not forming tight aggregated groups compared to the blue boxed region. We suspect that these lymphocytes are collecting in this region but are not forming a TLS due to geospace’s constraint of the tissue, or perhaps that this region represents the edge of a TLS, which is not present due to the relative thinness of the histology image. TLS is an ectopic lymphoid formation within nonlymphoid tissue. TLS can additionally foster tumor antigen presentation and T cell activation and the presence of TLS has been associated with improved response to cancer immunotherapy and prolonged patient survival [41-43]. Similar analysis was performed on the cutaneous malignant melanoma data[18] using canonical cell type marker genes **(Supplementary Fig. 11)**.

**Fig. 4.**
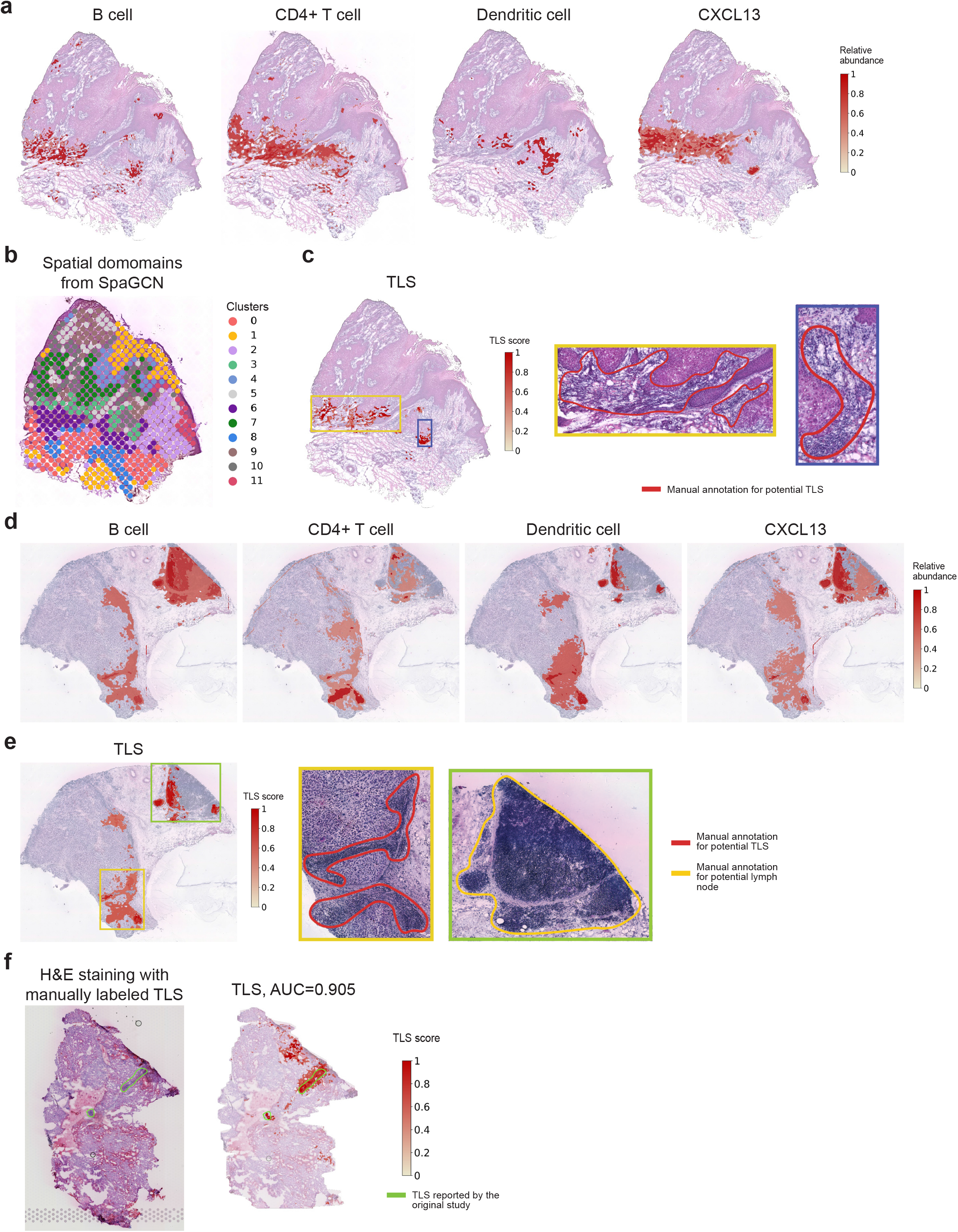
TESLA identifies lymphoid cell types and TLS with pixel-level resolution in the CSCC and melanoma samples. **a**, The distribution of B cells, CD4+ T cells, Dendritic cells and chemokine CXCL13 at pixel-level resolution characterized by TESLA in the CSCC tissue section. **b**, Clustering of spots of the CSCC tissue section using SpaGCN. **c**, TLS scores calculated by TESLA for the cutaneous CSCC tissue section. The blue box marks a highly possible TLS while the green box marks a region with aggregated lymphocytes but not forming a TLS. **d**, The distribution of B cells, CD4+ T cells, Dendritic cells, and chemokine CXCL13 at pixel-level resolution characterized by TESLA in the melanoma tissue section. **e**, TLS scores calculated by TESLA for the melanoma tissue section. The yellow and green boxes mark two potential TLSs and the blue box marks a lymphoid node apart from the tumor region. **f**, H&E staining and TLS scores calculated by TESLA for the clear cell renal cell carcinoma (ccRCC) primary tumor data. The yellow line outlines the true TLS annotated by pathologists, which was further validated by triple immunofluorescence labeling (CD20/CD3/PD-1) on an immediately adjacent slide for frozen sample and CD3/CD20 labeling by immunohistochemistry for FFPE sample.

We further validated the cell type distribution by TESLA in **Fig. 4a**,**d** by performing cell type deconvolution analysis using RCTD with annotated single-cell RNA-seq data from Ji et al. [17] and Tirosh et al. [19] as the references for the CSCC data and the melanoma data, respectively. The deconvolution results revealed similar distribution and density for B and T cells as predicted by TESLA **(Supplementary Figs. 12 and 13)**. These results also demonstrate that by utilizing known marker information, TESLA can infer cell subtypes without the reliance on single-cell reference, which is advantageous compared to reference-based deconvolution methods [24].

**Fig. 4e** shows TLS density predicted by TESLA. Aggregations of lymphocytes are observed at the bottom and the right side of the melanoma region, indicating strong immune activity in the yellow and blue boxes, where we observed co-localization of B cells, CD4+ T cells, and dendritic cells with a high expression of CXCL13 **(Fig. 4d)**. Since the top-region in the green box is not adjacent to tumor region but with aggregated lymphocytes, it is annotated as a lymph node instead of a TLS. These results highlight the importance of maintaining the spatial relationships of gene expression in heterogeneous tissue as the gene signature of TLS versus normal reactive lymph nodes are highly similar, reflecting the formation of lymph-node like architecture of TLS’s associated with tumors. By identifying the neoplastic tissue core and edge **(Fig. 3f)**, together with the TLS signature **(Fig. 4e)**, TESLA can segment tumor associated TLS versus adjacent non-neoplastic lymph node, which would have been lost in dissociated or homogenized tissue analyses such as single-cell transcriptomics. The presence of TLS was further validated by the co-location of follicular helper T cells and B cells **(Supplementary Fig. 14)**.

In addition to the CSCC and cutaneous malignant melanoma data, we also analyzed a dataset [44] that has manual annotation and immunostaining validated TLS information. This dataset was obtained from human clear cell renal cell carcinoma (ccRCC) primary tumors, where the TLSs were manually annotated by pathologists and further validated by triple immunofluorescence labeling (CD20/CD3/PD-1) on an immediately adjacent slide for frozen samples and CD3/CD20 labeling by immunohistochemistry for Formalin-Fixed Paraffin-Embedded (FFPE) samples. We performed TLS detection using TESLA on a tissue section in this dataset. As shown in **Figure 4f**, the TLS scores agree well with the two validated TLS regions. We then treated the TLS score, ranging from 0 to 1, as probability of being a TLS and calculated the pixel-wise AUC score using pathologist’s annotation provided in the original study as ground truth. TESLA achieved an AUC of 0.905, which further demonstrates TESLA’s ability to correctly locate the TLSs.

### Inferring cancer subtypes in breast cancer

Next, we show that integration of gene expression and histology information can help infer cancer subtypes. We first analyzed a HER2+, ER+ and PR-breast cancer dataset generated by 10x Genomics [20]. **Fig. 5a** summarized the cancer subtypes distribution, which is concordant with the diagnosis. We next analyzed an annotated HER2+ human breast cancer dataset from Andersson et al. [45]. We selected three patients H, G, and B and analyzed one tissue section from each patient. All three patients are HER2 positive and ER negative, and only Patient B is PgR positive while the other two are PgR negative. Based on the expression of basal markers such as *KRT5, KRT7, KRT14*, and *KRT18*, and breast cancer markers previously reported, e.g., *ERBB2, GATA3, PIP*, and *SCGB2A2* (**Supplementary Fig. 15**), TESLA detected the tumor region, demonstrating concordance with manual annotation from the original study **(Fig. 5b)**.

**Fig. 5.**
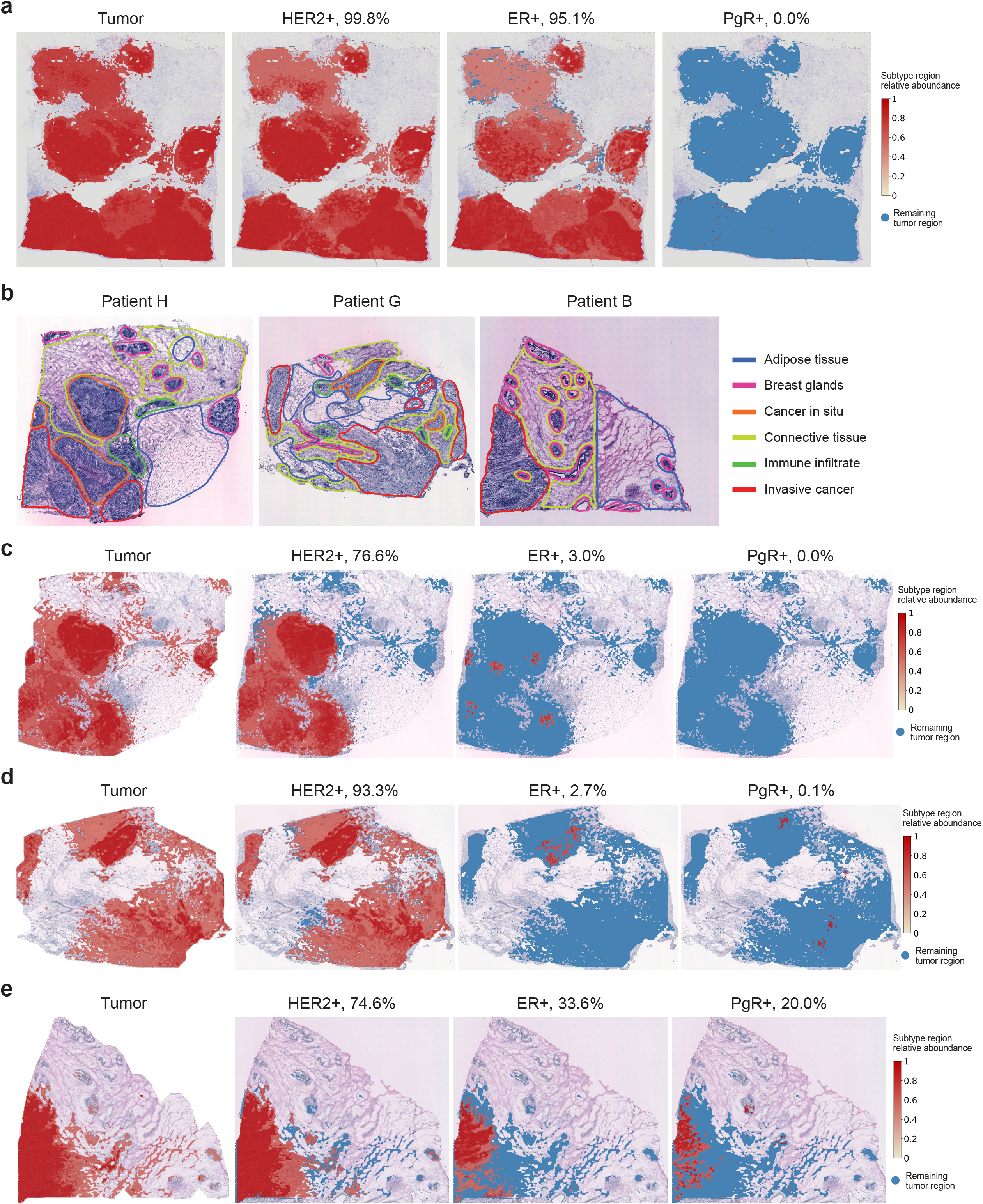
TESLA detects subtypes of breast cancer samples. **a**, Tumor region and tumor subtype regions (HER2+, ER+, and PgR+) detected by TESLA in the HER2+, ER+ and PR-breast cancer dataset from 10x Genomics. **b**, Histology images with pathologists’ manual annotation from the original HER2+ breast cancer study. **b-d**, Tumor region and tumor subtype regions (HER2+, ER+ and PgR+) detected by TESLA in the three breast cancer tissue sections from patients H, G and B in the HER2+ breast cancer study.

Next, based on specific marker gene expression, we detected the HER2+, ER+ and PgR+ regions within the tumor for each tissue section. **Fig. 5b-d** summarized the cancer subtypes distribution and the percentages of area for HER2, ER, and PgR for the three tissue sections. We observed high expression of *ERBB2*, which encodes HER2, in the tumors of all 3 patients, presented in 76%, 94%, and 76% of tumor cells, respectively, for patients H, G, and B, confirming the diagnosis of HER2+ cancers. In addition, we detected the expression of ER and PgR in patient B **(Fig. 5d)**, and low-level expression (0%-3%) of ER and PgR in the tumors of patients H and G, indicating the high degree of intratumoral heterogeneity in these tumors, owing to the super-resolution gene expression images generated by TESLA. For patient B, although it is diagnosed to be ER- and PR+, we detected 33.6% of the tumor cells to be ER+ and only 20.0% of the tumor cells to be PR+. We contacted the authors of the original paper and they claimed that the cancer subtype classifications were based on IHC (and PAM50) on a separate piece of the tumor. The low proportion of PR+ tumor cells and relatively high proportion of ER+ tumor cells may be due to the difference in hormone receptors between the piece used for the Visium experiment and the piece used for IHC. These results demonstrate the power of TESLA in quantitative evaluation of tumor subtypes and the additional information it provides that is beyond the simple positive or negative annotation of a tumor.

## Discussion

In this paper, we presented TESLA, a machine learning framework for multi-level tissue annotation on the histology image with pixel-level resolution in ST. By integrating information from high-resolution histology images, TESLA can impute gene expression at superpixels and fill in missing gene expression in tissue gaps. The increased gene expression resolution makes it possible to treat gene expression data as images, which enabled the integration with histological features for joint tissue segmentation and annotation of different cell types directly on the histology image with pixel-level resolution. Additionally, TESLA can detect unique structures of tumor immune microenvironment such as TLSs, separate a tumor into core and edge to examine their cellular compositions, expression features, and molecular processes. Principally, TESLA can distinguish lymphoid aggregates, immature and mature TLSs, if appropriate marker genes are provided by the users, but this will also depend on the quality of the ST data such as sequence coverage, the average number of genes detected in each spot, and dropout rate. TESLA has been evaluated on six cancer datasets, including cutaneous squamous cell carcinoma of the skin [17], cutaneous malignant melanoma[18], three breast cancer datasets [20, 21, 26], and clear cell renal cell carcinoma [44]. We further conducted a benchmark evaluation based on data generated from Stereo-seq [46] and showed that with TESLA enhanced gene expression, we can recover most of the differentially expressed genes that are only detectable in the original single-cell resolution Stereo-seq data **(Supplementary Note 3)**. Our results consistently showed that TESLA can generate high-quality super-resolution gene expression images, which facilitated the downstream multi-level tissue annotation.

Hematoxylin and eosin-stained histology images are routinely used in clinics for disease diagnosis. Utilizing manual annotation of histology images provided by pathologists, supervised methods have been developed to computationally annotate tissues; for example, Saltz et al. [47] developed a deep learning model focusing on detecting TIL using histology images in The Cancer Genome Atlas Program. However, these supervised methods require large, well-labelled training data in which expert pathologists are demanded to review a large set of images and mark detailed regions of lymphocytes and necrosis. Although pathologists’ annotation is highly effective for disease diagnosis, it is qualitative and prone to both inter- and intra-observer variability, particularly when quantifying or characterizing feature-rich phenomena such as tumor-associated lymphocytic infiltrates. Additionally, existing supervised methods only utilize histology information, which limit their usefulness in studying detailed structure in TME as histology alone cannot reveal subtypes of lymphocytes but can be distinguished using gene expression.

The cell type annotation in TESLA shares similarity with traditional clustering-based cell type annotation, but TESLA has several advantages. First, TESLA’s annotation has much higher resolution than annotation obtained from spot-level gene expression clustering. Second, in traditional clustering, each spot is assigned to a single cell type. However, for spatial barcoding-based ST data, e.g., Spatial Transcriptomics and 10x Visium, each spot may contain multiple cells. Although deconvolution-based methods such as RCTD [24] can infer cell type proportions within each spot, they cannot tell the cell identity for each cell within the spot. By contrast, through leveraging high-resolution histology image information, TESLA’s annotation can provide cell type enrichment information with pixel-level resolution. Additionally, deconvolution-based methods require an annotated single-cell reference [24, 48-50], which would limit their applications when single-cell references are not available.

In this paper, we only showed examples of combining histology and gene expression for cell type annotation, but it is also possible to incorporate immunofluorescence images for cell-type-specific proteins to infer targeted cell types using TESLA. Although immunofluorescence images can be used directly to infer cell identity for each cell, artifact in the images may render the results less reliable. When analyzing the CD3ε immunofluorescence data, we noticed the light in the original CD3ε immunofluorescence image is not flat across the tissue area, leading to brighter staining in the upper part and darker staining in the lower part of the tissue **(Fig. 2c)**. This uneven illumination results in an inaccurate estimation of CD3E abundance. By combining the gene image for *CD3E* and the CD3E immunofluorescence image into a 2-channel image as input, TESLA inferred the distribution of T cells. As shown in **Supplementary Fig. 16**, the upper edge now has lower degree of T cell enrichment, suggesting that combining gene expression and immunofluorescence image can correct immunofluorescence image artifact.

Although we illustrated the application of TESLA in cancer, the framework is generic and can be applied to other medical conditions as long as high-resolution histology images are available. We also demonstrate that TESLA is computationally fast and memory efficient compared to BayesSpace **(Supplementary Note 4)**. With the increasing popularity of ST in biomedical research, we expect TESLA will be an attractive tool for ST data analysis. Results from TESLA will enable researchers to increase gene expression resolution and annotate their tissues of interest with high confidence.

## Acknowledgements

This work was supported by the following grants: R01GM125301 (to M.L.) and P01AG066597 (to E.B.L. and M.L.). L.W. was supported in part by the start-up research fund and the institutional research grant (IRG) awards provided by U.T. MD Anderson Cancer Center, the Andrew Sabin Family Fellowship provided by the Andrew Sabin Family Foundation, and the RP200385 award provided by Cancer Prevention & Research Institute of Texas (CPRIT). We thank Dr. Kim Thrane for sharing the melanoma histology image data.

## Author contributions

This study was conceived of and led by M.L.. J.H. designed the model and algorithm. J.H. implemented the TESLA software and led the data analysis with input from M.L., L.W., E.B.L., H.K. and K.C.. K.C. performed cell type deconvolution analysis. E.B.L. examined the histology images. L.W. and H.K. provided marker genes for cancer and interpretated the results. L.W. provided guidance for the tumor TME analysis. J.H., M.L., and L.W. wrote the paper with feedback from all other coauthors.

## Methods

### Data preprocessing

TESLA takes spatial gene expression and histology image data as input. The spatial gene expression data contain an *N* × *G* matrix of unique molecular identifier (UMI) counts with *N* spots and *G* genes, along with the (*x, y*) 2-dimensional spatial coordinates of each spot. The gene expression values in each spot are normalized such that the UMI count for each gene in a given spot is divided by the total UMI count across all genes in that spot, multiplied by 10,000, and then transformed to a natural log scale.

### Super-resolution gene expression image generation

After preprocessing, TESLA extracts the measured tissue region from the histology image and generates a super-resolution gene expression image for each gene in the tissue using histological, spatial location, and the original spot-level gene expression data. The super-resolution gene expression generation step in TESLA is based on two assumptions: 1) spots that are physically close have similar gene expression, and 2) the expression patterns for spatially variable genes are correlated with histology image features. Therefore, this enhancement step works better for genes whose expression patterns have some spatial variability as compared to non-spatially variable genes. Below we describe each step in super-resolution gene expression generation in detail.

#### Superpixel generation

TESLA first detects the contour of the captured tissue area in the histology image. When the tissue region can be easily separated from the histology image background, TESLA uses the Canny-edge-detection algorithm implemented in the python package “opencv” to draw the contour. When the histology image contains contaminated stains or tissue fragments, the Canny-edge-detection algorithm may result in an inaccurate contour. In this case, TESLA detects the boundary of the measured spots and uses the enlarged boundary as the tissue contour. After contour detection, TESLA separates the tissue region inside the contour into equal-sized squares, referred to as superpixels. The size of the superpixels can be adjusted depending on the dataset and is usually much smaller than spots captured by next generation sequencing-based ST technologies. The default size of a superpixel is (50 × 50) pixels, but can be adjusted by user depending on the diameter size of each spot in the ST data. As a comparison, the diameter is ∼350 pixels for a Spatial Transcriptomics spot and ∼200 pixels for a 10x Visium spot **(Supplementary Table 1)**.

#### Superpixel gene expression imputation

We aim to impute the gene expression at each superpixel using observed spot-level gene expression. The gene expression of a superpixel is expected to be similar to that of its neighboring spots. Thus, to impute the gene expression at a given superpixel *ν*, TESLA detects its top 10 nearest neighboring measured spots based on the Euclidean distance metric described in SpaGCN[25], which considers the similarity between the superpixel *ν* and a measured spot with respect to both physical location and histological features. We first extract the approximate region of each superpixel on the histology image and calculate the mean color value for the RGB channels, (*r*_*ν*,_*g*_*ν*,_*b*_*ν*_), of all pixels that fall in that region. Next, a weighted sum of the RGB value is calculated to represent the histology image features,

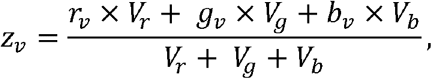

where *V*_*r*_ = Variance(*r*_*ν*_), *V*_*g*_= Variance(*g*_*ν*_), and *V*_*b*_ = Variance(*b*_*ν*_) for all *ν* ∈ *V*. Then, *z*_*ν*_ is rescaled to the same scale as *x*_*ν*_ and *y* _*ν*_ as

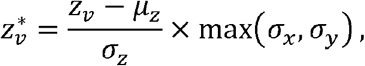

where *μ*_*Z*_ is the mean of, *z*_*ν*_, *σ*_*x*_, *σ*_*y*_, *σ*_*z*_ are the standard deviations of *x*_*ν*,_ *y*_*ν*_ and *z*_*ν*_ respectively. The 3D coordinates for all the measured spots are derived using the same approach, and the Euclidean distance between superpixel *ν* and a measured spot *u* is calculated as

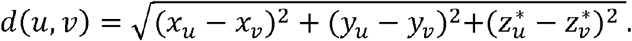

For each superpixel *ν*, TESLA detects its top 10 nearest neighboring measured spots, denoted as set *N*_*ν*_. Then, the gene expression value of the superpixel *ν* is imputed using a weighted sum of the neighboring spots’ gene expression values. The weight for a given neighboring spot *u*_*i*_ *∈ N*_*ν*_ is negatively associated with the distance from that spot to the superpixel and is defined as

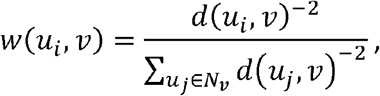

where *d*(*u*_*i*,_ *ν*) is the Euclidean distance calculated using the approach described in SpaGCN. For a given gene *g*, its imputed expression value for superpixel *ν* is calculated as

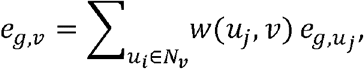

where 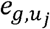 is the measured spot-level gene expression for gene *g* at spot *u*_*i*_. For each pixel within the superpixel, its expression is imputed as the gene expression at the corresponding superpixel.

After imputation, TESLA generates a super-resolution gene expression image for each gene in the tissue with tissue gaps imputed as well. We note that the center-to-center distance between two adjacent spots in the current 10x Visium platform is 100 μm, leaving a large portion of the tissue unmeasured for gene expression. Although BayesSpace can enhance the gene expression resolution within measured spots, regions that are not covered by spots are still left blank.

### Multi-level tissue annotation

TESLA can annotate a tissue section at multiple levels by integrating different super-resolution gene expression images together with a histology image for joint segmentation using a convolutional neural network approach. For example, using integrated cell-type-specific marker gene images and the histology image as input, TESLA can reveal cell type distributions at the same pixel-level resolution as the histology. Additionally, TESLA can identify tumor regions if the tumor marker gene images are used as input for joint segmentation. The joint consideration of tumor marker genes and histology images enables the detection of intra tumor heterogeneity that might be missed by traditional pathology-based tumor diagnosis. For simplicity, we use cell type annotation to demonstrate the annotation method.

#### Generation of meta gene image

The joint segmentation takes both gene expression and histology images as input. A histology image has 3 color channels representing red (R), green (G), and blue (B) color intensities, while the gene expression are represented by *K* channels for a cell type with *K* marker genes. If the channels are weighted equally, the gene expression would dominate the joint segmentation results when *K* is larger than 3, whereas the histology image would dominate when *K* is less than 3. Therefore, it is important to find a balance between the histology and gene expression. Since the number of marker genes varies across cell types, we combine the *K* markers into one meta marker gene, which can be represented by one color channel, to make our method robust across different cell types.

Since not all marker genes are expressed, an ideal meta gene should preserve the expression pattern for at least a subset of the marker genes. Given *K* marker genes and a pre-specified number *k*, for any superpixel *i*, we first rank all markers’ relative expression in descending order as {*e*_1*i*_, *e*_2,*i*_, *e*_3,*i*_,… *e*_*K,i*_},. Next, we select the top *k* expression values and calculate the meta gene’s relative expression at superpixel *i* as:

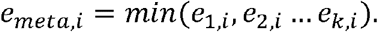

At a given superpixel, if only the top *k* − 1 or fewer marker genes are expressed and the *k*^*th*^ marker has zero expression, the meta gene will have an estimated expression value of 0 at that superpixel. This step ensures that expression patterns that are present in less than *k* genes are not included in the meta gene, thus avoiding generating patterns that are less representative. Defining the meta gene’s expression in this way guarantees that its expression level is high at a given superpixel if and only if all top *k* markers are highly expressed at that superpixel. In our analyses, we set *k* equal to 1 to capture the expression patterns of all marker genes.

#### Tissue segmentation using convolutional neural network

Next, we convert the RGB channels from the histology image into one gray channel. For the meta gene image, we use the gene expression at each superpixel to represent the expression for each pixel that resides within the superpixel. Since the gray and meta gene channels are the same size, we can simply stack them to create a 2-channel image, where each pixel value is normalized to [0, 1] and fed into a convolutional neural network for unsupervised segmentation[51] **(Fig. 6)**. Let *M* be the number of pixels in the 2-channel image denoted by 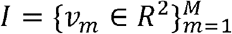. A p-dimensional feature map 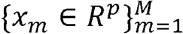 is computed from 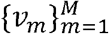 through two convolutional components, each of which consists of a 2-dimensional convolution, a ReLU activation function, and a batch normalization function. Subsequently, a response map 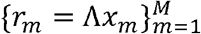 is obtained by applying a linear classifier on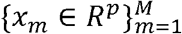, where Λ ∈ *R* ^*p*×*p*^ represents the weights in the convolutional layer. The response map is then normalized to 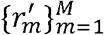, which has zero mean and unit variance. Finally, the initial cluster label *c*_*m*_ for each pixel is obtained by selecting the cluster that has the maximum value in 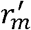. To initialize the neural network, we start with a large initial number of clusters (default *p* = 100).

**Fig. 6.**
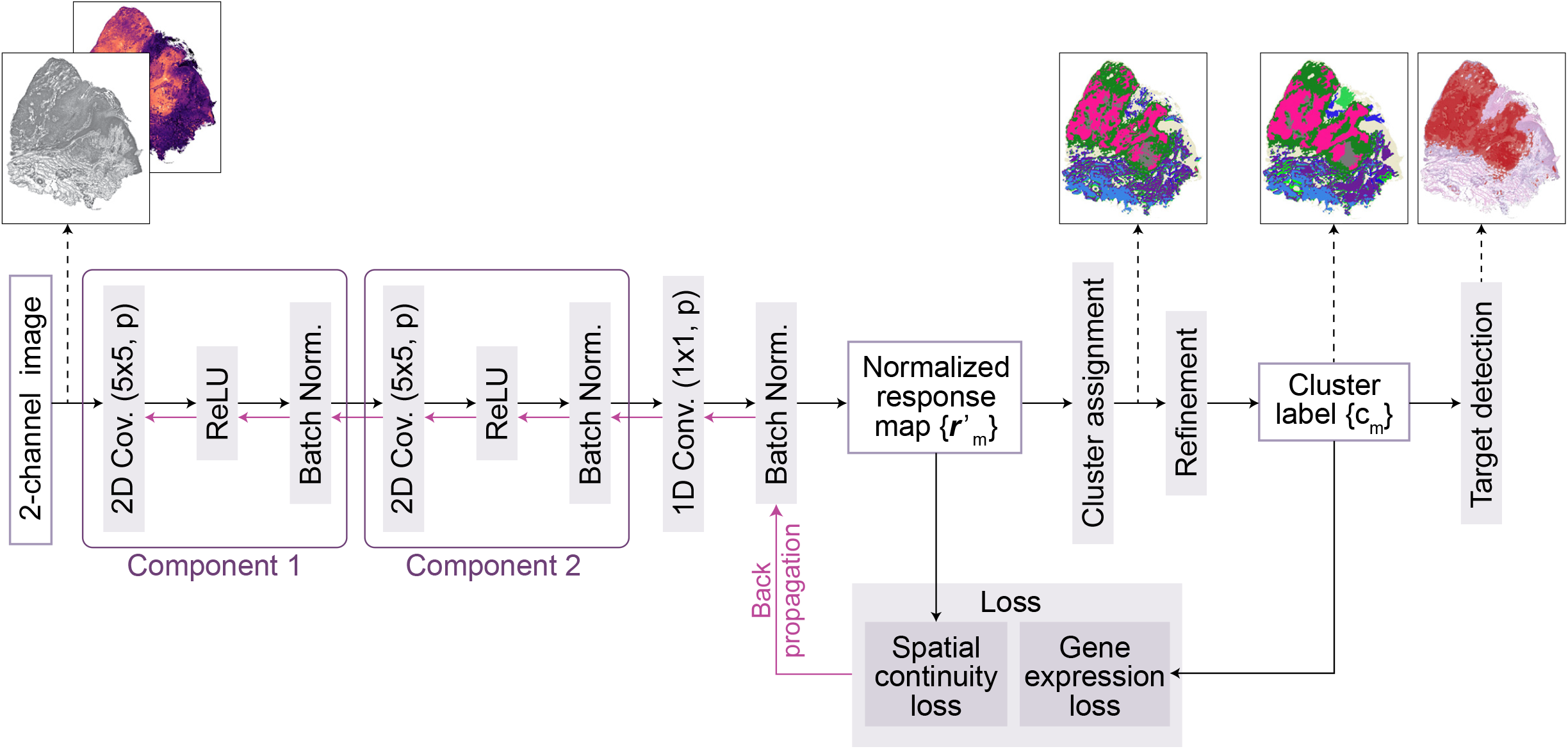
Architecture of the CNN for image segmentation in TESLA. A 2-channel image is fed into the CNN to extract deep features through two convolutional components, each of which consists of a 2-dimensional convolution (5×5, *p*), a ReLU activation function, and a batch normalization function. Subsequently, the response vectors of the features in *p*-dimensional cluster space are calculated through a one-dimensional convolutional laye (1×1, *p*), and normalized to 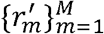 across the axes of the cluster space using a batch normalization function. The normalized response map 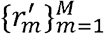 is used to calculate the spatial continuity loss. Further, raw cluster labels are determined by assigning the cluster to 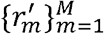 using an argmax function. Next, the raw cluster labels are refined to 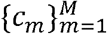 and used as pseudo labels to compute the gene expression loss. Finally, the spatial continuity loss as well as the gene expression loss are combined and backpropagated. After training, this CNN can segment input image into different clusters, and TESLA next identifies the clusters corresponding to the target region.

The loss function *L* consists of a constraint on spatial continuity and a constraint on gene expression similarity, defined as follows,

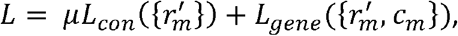

where *μ* represents the weight for balancing the two constraints (*μ* = 5 as default). Following Kim et al.[28], we utilize the L1-norm of horizontal and vertical differences of the response map 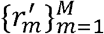 as a spatial constraint, which is defined as

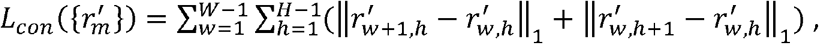

where *W* and *H* represent the width and height of the 2-channel image, and 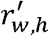, represents the pixel value at (*w,h*) in the response map 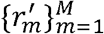. By applying this spatial continuity loss, an excessive number of clusters due to complicated patterns can be suppressed. The following cross entropy loss between 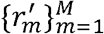 and 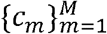 is calculated as the constraint on gene expression similarity,

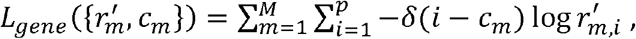

where *δ*(·) is an indicator function. This loss ensures that pixels assigned to the same cluster have similar meta gene expression.

In the iterative update procedure, cluster memberships for spatially close pixels or pixels with similar meta gene expression values are merged by considering spatial continuity and gene expression similarity. This process leads to the reduction in the number of unique clusters *p*^′^. The normalized response map vectors and cluster assignments are updated iteratively as follows. Since some clusters may have no pixels assigned to them, the number of unique clusters can be a number *p*^′^ ranging from 1 to *p*. During each training epoch, TESLA has a cluster refinement step to merge minor clusters to its neighboring major clusters. First, clusters containing less than 1000 (default) pixels will be selected as minor clusters. Next, for each pixel in a given minor cluster, its neighboring pixels, defined as pixels located within a circle of radius 3 centered at that pixel, are collected into a set. Based on the cluster identities of all neighboring pixels of the pixels in that minor cluster, TESLA will detect the major cluster which has the most pixels in the neighboring set, referred to as the neighboring major cluster. TESLA then merges the minor cluster with its neighboring major cluster, which increases the integrity of the segmentation. Due to the cluster refinement step, the number of clusters decreases during the training process. The model will stop training when the number of clusters reaches a pre-defined threshold (default is 30). We tested the value of this threshold from 10 to 50 and found that it does not significantly affect the target cell type detection. We use 30 as default to allow the algorithm to assign some tissue edge and small contaminated regions as small clusters.

#### Target cell type detection

After obtaining the cluster assignments from segmentation, TESLA identifies which cluster(s) are enriched for a specified target cell type. Since the segmentation output has the same resolution as the meta gene image, TESLA can calculate the average meta gene relative expression across all pixels within each cluster and sort the clusters in descending order of this value as {*ē*_*meta,cluster*1_, *ē*_*meta,cluster*2_,…, *ē*_*meta,clusterp*_}. Next, clusters whose average meta gene relative expression is greater than *t*× *ē* _*meta,cluster*1_ are considered as clusters that are enriched for the target cell type. We find that *t* = 1/2 generally generates the best performance in practice. Finally, each cluster is annotated on the histology image with color corresponding to the average meta gene expression level. This framework is flexible, where users can customize their own gene list for a cell type of interest; for example, by changing the cell type marker genes to tumor marker genes **(Supplementary Table 4)**, one can detect the tumor region in the tissue.

### Intra-tumor heterogeneity analysis

With tumor region annotation, we can further draw the leading edge and separate the tumor into edge and core, which allows us to study intra-tumor heterogeneity. TESLA first detects the boundary of the tumor region using the Canny-edge-detection algorithm implemented in the python package “opencv”. Tumor edge is defined as the area that is within 200 μm interface of tumor and adjacent to non-malignant tissue and the tumor core is defined as the proximal tumor area within the tumor edge[28]. After the tumor edge and core are identified, we can perform differential expression analysis to detect genes enriched in the edge and the core, which will help in understanding the heterogeneity inside the tumor. We performed DE analysis between pixels in the tumor core and edge using Wilcoxon rank-sum test. Genes with a false discovery rate (FDR) adjusted p-value <0.05 are first selected. To ensure only genes with enriched expression patterns in the core/edge region are selected, we further require a gene to meet the following three criteria: 1) mean log (gene expression) >0.1; 2) in/out group expression fraction ratio >=1, where “in group” refers to spots in the target region (e.g., tumor edge or tumor core), and “out group” refers to the remaining spots, “in group expression fraction” is the percentage of spots in the “in group” where the gene is expressed, and “out group expression fraction” is the percentage of spots in the “out group” where the gene is expressed; 3) in group expression fraction >0.5; 4) fold change of in group gene expression/out group gene expression >1.2.

### Detection of tertiary lymphoid structure

Previous studies have suggested that TLS formation results from a complex interplay between B cells, CD4+ T cells, and dendritic cells, with reciprocal signaling between these cells mediated by chemokine CXCL13[10]. TESLA detects the potential location of TLS based on the colocalization of B cells, CD4+ T cells, dendritic cells, and chemokine CXCL13 (marker genes in **Supplementary Table 4**). For any pixel *i*, TESLA first calculates the molecular abundance of each cell type as the mean expression across all pixels for the corresponding meta gene of that cell type. TELSA then ranks the molecular abundances of these cell types and CXCL13 in descending order as {*b*_1,*i*_, *b*_2,*i*_, *b*_3,*i*,_ *b*_4,*i*,_}. Next, the TLS score is calculated as the minimum value of the top three abundances:

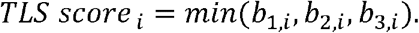

This minimum pooling from the top three values ensures that a pixel has a high TLS score if and only if at least three out of the four abundances are high. We do not perform minimum pooling using all four values because we expect the enrichment of B cells, CD4+ T cells, and dendritic cells, and CXCL13 may be closely related to TLS but not perfectly overlap with the other three cell types. Next, we annotate the whole tissue with color corresponding to the TLS score, where dense regions embedded inside the tumor with high TLS scores are identified as TLSs.

### Comparing image similarity using SSIM

The Structural SIMilarity (SSIM) index is a method for measuring the similarity between two images. The SSIM between two images *x* and *y* of the same size is calculated as

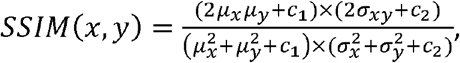

where (*μ*_*x*_, *μ*_*y*_) and (*σ*_*x*_, *σ*_*y*_) are the means and standard deviations of *x* and *y, σ*_*x,y*_ is the covariance of *x* and *y, c*_1_ (0.01 × *L)*^2^ and *c*_2_ = (0.03 × *L*)^2^ are two variables used to stabilize the division, and *L* is the dynamic range of the pixel-values. When calculating the SSIM between protein’s immunofluorescence staining image and the super-resolution gene image, the protein staining intensity and the super-resolution gene expression are scaled to the range (0, 255).

## Data availability

We analyzed eight spatial transcriptomics datasets and two scRNA-seq datasets. Publicly available data were acquired from the following websites or accession numbers: (1) human Invasive ductal carcinoma 10x Visium data (https://support.10xgenomics.com/spatial-gene-expression/datasets/1.2.0/V1_Human_Invasive_Ductal_Carcinoma); (2) human cutaneous squamous cell carcinoma 10x Visium data (GSE144240); (3) human cutaneous squamous cell carcinoma scRNA-seq data from 10X Chromium 3’ v2 (GSE144240); (4) human cutaneous malignant melanoma ST data (https://www.spatialresearch.org/resources-published-datasets/doi-10-1158-0008-5472-can-18-0747/); (5) human melanoma tumor scRNA-seq data from Smart-Seq2 (GSE72056); (6) human HER2+, ER+ and PR-breast cancer dataset from 10x Genomics (https://support.10xgenomics.com/spatial-gene-expression/datasets/1.1.0/V1_Breast_Cancer_Block_A_Section_1); (7) human HER2-positive breast tumor ST data (https://github.com/almaan/her2st); (8)Mouse posterior brain (sagittal) 10x Visium data (https://support.10xgenomics.com/spatial-gene-expression/datasets/); (9) Mouse kidney (coronal) 10x Visium data (https://support.10xgenomics.com/spatial-gene-expression/datasets/); (10) Human clear cell renal cell carcinoma primary tumors 10x Visium data (GSE175540).

Details of the datasets analyzed in this paper were described in **Supplementary Table 1**.

## Software availability

An open-source implementation of the TESLA algorithm in Python can be downloaded from https://github.com/jianhuupenn/TESLA

## Life sciences reporting summary

Further information on experimental design is available in the Life Sciences Reporting Summary.

