## Supplement for "Deciphering tumor ecosystems at super-resolution from spatial transcriptomics with TESLA"

**Supplementary Table 1. Datasets analyzed in this paper.**

| **Species** | **Tissue** | **Data source** | **Dataset dimensions** | **Protocol** | **Spot diameter by pixels** |
| --- | --- | --- | --- | --- | --- |
| Human | Invasive ductal carcinoma | 10x Genomics  (https://support.10xgenomics.com/spatial-gene-expression/datasets/1.2.0/V1_Human_Invasive_Ductal_Carcinoma)[1] | 4,727 spots  36,601 genes | 10x Visium | 172 pixels |
| Human | Cutaneous squamous cell carcinoma | Ji *et al*.[2]  GSE144240 | 646 spots  17,344 genes | 10x Visium | 200 pixels |
| Human | Cutaneous squamous cell carcinoma | Ji *et al*.[2]  GSE144240 | 6,824 cells  32,738 genes | 10X Chromium 3' v2 | NA |
| Human | Cutaneous malignant melanoma | Thrane *et al*.[3]  (https://www.spatialresearch.org/resources-published-datasets/doi-10-1158-0008-5472-can-18-0747/) | 293 spots  16,148 genes | Spatial Transcriptomics | 350 pixels |
| Human | Melanoma tumor | Tirosh *et al*.[4] [https://science.sciencemag.org/content/352/6282/189]  GSE72056 | 4,139 cells  23,686 genes | Smart-Seq2 | NA |
| Human | HER2+, ER+ and PR- breast cancer | 10x Genomics  (https://support.10xgenomics.com/spatial-gene-expression/datasets/1.1.0/V1_Breast_Cancer_Block_A_Section_1)[5] | 3,798 spots  36,601 genes | 10x Visium | 172 pixels |
| Human | HER2-positive breast tumor | Andersson *et al*.[6]  (https://github.com/almaan/her2st) | 295 spots  15,109 genes | Spatial Transcriptomics | 146 pixels |
| Mouse | Posterior brain (sagittal) | 10x Genomics  (https://support.10xgenomics.com/spatial-gene-expression/datasets/)[7] | 3,355 spots  32,285 genes | 10x Visium | 80 pixels |
| Mouse | Kidney (coronal) | 10x Genomics  (https://support.10xgenomics.com/spatial-gene-expression/datasets/)[8] | 1,438spots  32,285genes | 10x Visium | 80 pixels |
| Human | Clear cell renal cell carcinoma primary tumors | Meylan *et al*.[9]  GSE175540 | 4,359spots  36,945genes | 10x Visium | 24 pixels |

**Supplementary Table 2. Overlaps with curated gene sets included in the Molecular Signature Database by performing gene set enrichment analysis using region-specific DEGs in the cutaneous squamous carcinoma dataset.**

| **Tumor core enriched genes, n=300** | | | | |
| --- | --- | --- | --- | --- |
| Name | Description | n | FDR q-value | Genes |
| REACTOME_METABOLISM_OF_LIPIDS | Metabolism of lipids | 26 | 2.46e-07 | AGPAT3, LPIN3, TAZ, PLD1, PNPLA8, LCLAT1, PISD, PLBD1, PITPNM3, MTM1, PI4K2B, HADH, MMUT, PCCB, MECR, MCAT, DECR2, ELOVL6, ALOX12B, FDXR, TBL1X, MTF1, NFYC, HSD17B1, DHRS7B, MED21 |
| REACTOME_GLYCEROPHOSPHOLIPID_BIOSYNTHESIS | Glycerophospholipid biosynthesis | 9 | 0.000589 | AGPAT3, LPIN3, TAZ, PLD1, PNPLA8, LCLAT1, PISD, PLBD1, PITPNM3 |
| REACTOME_PHOSPHOLIPID_METABOLISMREACTOME_PHOSPHOLIPID_METABOLISM | Phospholipid metabolism | 11 | 0.000589 | AGPAT3, LPIN3, TAZ, PLD1, PNPLA8, LCLAT1, PISD, PLBD1, PITPNM3, MTM1, PI4K2B |
| REACTOME_MITOCHONDRIAL_FATTY_ACID_BETA_OXIDATION | Mitochondrial Fatty Acid Beta-Oxidation | 5 | 0.00463 | HADH, MMUT, PCCB, MECR, MCATHADH, MMUT, PCCB, MECR, MCAT |
| REACTOME_VESICLE_MEDIATED_TRANSPORT | Vesicle-mediated transport | 18 | 0.00475 | AGPAT3, TUBB4A, KIF23, RAB8A, KIF20B, NBAS, BET1L, SYS1, EPGN, VPS37A, DENND2C, DENND1B, MON1A, EPS15L1, AP1M2, COPS7A, FCHO2, EXOC3 |
| REACTOME_SIGNALING_BY_RETINOIC_ACID | Signaling by Retinoic Acid | 5 | 0.00704 | DHRS4, PDHB, PDK2, DHRS3, RDH14 |
| REACTOME_CELL_CYCLE_MITOTIC | Cell Cycle, Mitotic | 15 | 0.00768 | LPIN3, TUBB4A, KIF23, RAB8A, LIG1, FEN1, CENPF, SEH1L, ZWINT, DHFR, CDC23, ANAPC15, TFDP2, CKS1B, PPP2R3B |
| REACTOME_FATTY_ACID_METABOLISM | Fatty acid metabolism | 8 | 0.0155 | HADH, MMUT, PCCB, MECR, MCAT, DECR2, ELOVL6, ALOX12B |
| REACTOME_CELL_CYCLE | Cell Cycle | 16 | 0.0178 | LPIN3, TUBB4A, KIF23, RAB8A, LIG1, FEN1, CENPF, SEH1L, ZWINT, DHFR, CDC23, ANAPC15, TFDP2, CKS1B, PPP2R3B, MLH1 |
| REACTOME_METABOLISM_OF_AMINO_ACIDS_AND_DERIVATIVES | Metabolism of amino acids and derivatives | 11 | 0.0248 | PDHB, PXMP2, GSTZ1, CKB, ALDH9A1, BBOX1, ALDH4A1, PYCR3, SLC25A10, GPT2, ASPG |
| --- |  |  |  |  |
| **Tumor edge enriched genes, n=106** | | | | |
| Name | Description | n | FDR q-value | Genes |
| REACTOME_INNATE_IMMUNE_SYSTEM | Innate Immune System | 18 | 5.19e-07 | PTPN6, C2, FPR1, C3, PTAFR, FCGR1A, LILRB2, LAIR1, PTPRJ, FCER1G, MME, RHOF, BIN2, STK10, FCGR3A, FYN, DUSP4, CLU |
| HALLMARK_INTERFERON_GAMMA_RESPONSE | Genes up-regulated in response to IFNG GeneID=3458 | 9 | 1.8e-06 | PTPN6, IL2RB, IRF8, FPR1, FCGR1A, GBP4, ST3GAL5, CMPK2, ZNFX1PTPN6, IL2RB, IRF8, FPR1, FCGR1A, GBP4, ST3GAL5, CMPK2, ZNFX1 |
| REACTOME_NEUTROPHIL_DEGRANULATION | Neutrophil degranulation | 12 | 2.46e-06 | PTPN6, FPR1, C3, PTAFR, LILRB2, LAIR1, PTPRJ, FCER1G, MME, RHOF, BIN2, STK10 |
| REACTOME_ADAPTIVE_IMMUNE_SYSTEM | Adaptive Immune System | 14 | 1.23e-05 | PTPN6, CD3D, CD3E, FYB1, C3, AKT3, FCGR1A, LILRB2, LAIR1, PTPRJ, FCGR3A, FYN, LILRB4, TRIM39 |
| REACTOME_CYTOKINE_SIGNALING_IN_IMMUNE_SYSTEM | Cytokine Signaling in Immune system | 12 | 0.00011 | CCR1, PTPN6, IL2RB, IRF8, FPR1, PTAFR, AKT3, FCGR1A, PTPRJ, FYN, DUSP4, GBP4 |
| KEGG_CHEMOKINE_SIGNALING_PATHWAY | Chemokine signaling pathway | 7 | 0.000156 | CCR1, CXCL13, CX3CL1, CCR7, CXCL12, AKT3, CCL18 |
| PID_TCR_PATHWAY | TCR signaling in native CD4+ T cells | 5 | 0.000156 | PTPN6, CD3D, CD3E, FYB1, FYN |
| HALLMARK_INFLAMMATORY_RESPONSE | Genes defining inflammatory response | 7 | 0.000187 | IL2RB, FPR1, PTAFR, CX3CL1, CCR7, GPR183, RGS16 |

**Supplementary Table 3. Overlaps with curated gene sets included in the Molecular Signature Database by performing gene set enrichment analysis using region-specific DEGs in the cutaneous malignant melanoma dataset.**

| **Tumor core enriched genes, n=300** | | | | |
| --- | --- | --- | --- | --- |
| Name | Description | n | FDR q-value | Genes |
| REACTOME_SIGNALING_BY_RHO_GTPASES_MIRO_GTPASES_AND_RHOBTB3 | Signaling by Rho GTPases, Miro GTPases and RHOBTB3. | 23 | 6.96e-06 | STAM, CKAP4, AAAS, H2AC19, PTK2, DOCK7, FNBP1L, ARHGEF12, PLXNA1, BAIAP2, SCRIB, ARHGAP32, STARD8, ALDH3A2, LRRC1, EMD, BAIAP2L1, ANKFY1, PAFAH1B1, DVL3, TRAK2, NF2, CENPT |
| PID_MET_PATHWAY | Signaling events mediated by Hepatocyte Growth Factor Receptor (c-Met) | 8 | 9.39e-05 | RAB5A, PTK2, MET, RANBP10, RANBP9, BCAR1, PXN, EIF4EBP1 |
| REACTOME_SIGNALING_BY_MET | Signaling by MET | 8 | 9.39e-05 | STAM, USP8, PTK2, DOCK7, MET, RANBP10, RANBP9, ITGA3 |
| REACTOME_RHO_GTPASE_CYCLE | RHO GTPase cycle | 16 | 0.000132 | STAM, CKAP4, AAAS, DOCK7, FNBP1L, ARHGEF12, PLXNA1, BAIAP2, SCRIB, ARHGAP32, STARD8, ALDH3A2, LRRC1, EMD, BAIAP2L1, ANKFY1 |
| REACTOME_SIGNALING_BY_RECEPTOR_TYROSINE_KINASESREACTOME_SIGNALING_BY_RECEPTOR_TYROSINE_KINASES | Signaling by Receptor Tyrosine Kinases | 15 | 0.00158 | STAM, USP8, PTK2, DOCK7, BAIAP2, MET, RANBP10, RANBP9, BCAR1, PXN, ITGA3, ATP6V1C1, TRIB3, COL9A3, ID4 |
| REACTOME_MET_ACTIVATES_RAS_SIGNALING | MET activates RAS signaling | 3 | 0.0121 | MET, RANBP10, RANBP9 |
| REACTOME_SIGNALING_BY_WNT | Signaling by WNT | 10 | 0.026 | H2AC19, USP8, PSME3, AXIN1, SCRIB, DVL3, ASH2L, WLS, LGR4, SOX13 |
| --- |  |  |  |  |
| **Tumor edge enriched genes, n=155** | | | | |
| Name | Description | n | FDR q-value | Genes |
| HALLMARK_EPITHELIAL_MESENCHYMAL_TRANSITION | Genes defining epithelial-mesenchymal transition, as in wound healing, fibrosis and metastasis | 14 | 6.66e-11 | THY1, CAPG, VCAM1, SDC1, LUM, LOXL1, LOXL2, PCOLCE, FBLN1, BGN, IL32, TGFBI, FSTL1, SFRP4 |
| REACTOME_EXTRACELLULAR_MATRIX_ORGANIZATION | Extracellular matrix organization | 15 | 7.62e-10 | ITGAL, MMP9, VCAM1, SDC1, LUM, LOXL1, LOXL2, PCOLCE, FBLN1, BGN, PECAM1, VWF, COL6A1, LTBP2, LAMB2 |
| BIOCARTA_TCYTOTOXIC_PATHWAY | T Cytotoxic Cell Surface Molecules | 5 | 3.36e-07 | THY1, ITGAL, CD8A, CD2, CD3E |
| REACTOME_ADAPTIVE_IMMUNE_SYSTEM | Adaptive Immune System | 18 | 1.43e-06 | ITGAL, CD8A, HLA-DOA, CD3E, PRKCB, FYB1, VCAM1, CD22, C3, PLCG2, TAB2, FCGR1B, SLAMF7, LILRB2, LAIR1, BLK, ANAPC2, LAG3 |
| REACTOME_CYTOKINE_SIGNALING_IN_IMMUNE_SYSTEM | Cytokine Signaling in Immune system | 16 | 5.79e-06 | MMP9, IL4R, IL2RB, CCL19, IRF8, VCAM1, SDC1, IL32, TAB2, FCGR1B, IL10RA, CEBPD, LGALS9, TNFSF13B, CD27, SAMHD1 |
| REACTOME_IMMUNOREGULATORY_INTERACTIONS_BETWEEN_A_LYMPHOID_AND_A_NON_LYMPHOID_CELL | Immunoregulatory interactions between a Lymphoid and a non-Lymphoid cell | 9 | 1.35e-05 | ITGAL, CD8A, CD3E, VCAM1, CD22, C3, SLAMF7, LILRB2, LAIR1 |
| BIOCARTA_THELPER_PATHWAY | T Helper Cell Surface Molecules | 4 | 2.03e-05 | THY1, ITGAL, CD2, CD3E |
| REACTOME_SIGNALING_BY_INTERLEUKINS | Signaling by Interleukins | 11 | 0.000247 | MMP9, IL4R, IL2RB, CCL19, VCAM1, SDC1, IL32, TAB2, IL10RA, CEBPD, LGALS9 |
| PID_CD8_TCR_DOWNSTREAM_PATHWAY | Downstream signaling in native CD8+ T cells | 5 | 0.000649 | CD8A, CD3E, PRKCB, IL2RB, GZMB |
| KEGG_CYTOKINE_CYTOKINE_RECEPTOR_INTERACTION | Cytokine-cytokine receptor interaction | 8 | 0.00105 | IL4R, IL2RB, CCL19, IL10RA, TNFSF13B, CD27, CCL21, CXCL14 |
| HALLMARK_INTERFERON_GAMMA_RESPONSE | Genes up-regulated in response to IFNG | 7 | 0.00139 | IL4R, IL2RB, IRF8, VCAM1, SLAMF7, IL10RA, SAMHD1 |

**Supplementary Table 4. Marker genes used for cell type, tumor region and protein detection.**

| **Target region** | **Marker genes** |
| --- | --- |
| B cell | *CD19, CD79A, CD79B, MS4A1, CD22* |
| CD8+ T cell | *CD8A, CD8B* |
| Follicular helper T cells | *CD3E, CD3D, CD3G, CD4, PDCD1, CXCR5* |
| Dendritic cell | *CD1A, CD1B, CD1E, CLEC10A, CLIC2, WFDC21P* |
| CXCL13 | *CXCL13* |
| Melanoma | *MITF, CSPG4, MAGEA1, MLANA, TYR, SOX10* |
| Squamous cell carcinoma | *BUB1B, KIF1C, TOP2A, CD151, MMP10, PTHLH, FEZ1, IL24, KCNMA, INHBA, MAGEA4, NT5E, LAMC2, SLITRK6* |
| Breast Cancer | *ERBB2, CNN1, CDH1, KRT5, KRT7, KRT14, KRT18, CDNND1, GATA3, FOXA1, PIP, SCGB2A2* |
| HER2+ tumor subtype | *ERBB2* |
| ER+ tumor subtype | *ESR1* |
| PgR+ tumor subtype | *PGR* |

**Supplementary Table 5. Software compared with TESLA**.

| **Method** | **Version** | **URL** | **Reference** |
| --- | --- | --- | --- |
| BayesSpace | 1.0.0 | https://github.com/edward130603/BayesSpace | Zhao *et al*.[1] |
| SpaGCN | 1.2.0 | https://github.com/jianhuupenn/SpaGCN | Hu *et al*.[10] |
| RCTD | 1.2.0 | https://github.com/dmcable/RCTD | Cable *et al*.[11] |

**Supplementary Fig. 1.** Boxplot of Pearson correlations between the original gene expression and TESLA’s imputed gene expression for masked spots for top 2000 HVGs (n=2000 for all datasets). The goal of these hold-off experiments is to evaluate whether TESLA is able to recover the original gene expression. In this analysis, we masked each spots in the 10x Visium data and only used the remaining spots as known data to impute gene expression for that masked spot. The lower and upper hinges correspond to the first and third quartiles, and the center refers to the median value. The upper (lower) whiskers extend from the hinge to the largest (smallest) value no further (at most) than 1.5 × interquartile range from the hinge. Data beyond the end of the whiskers are plotted individually.

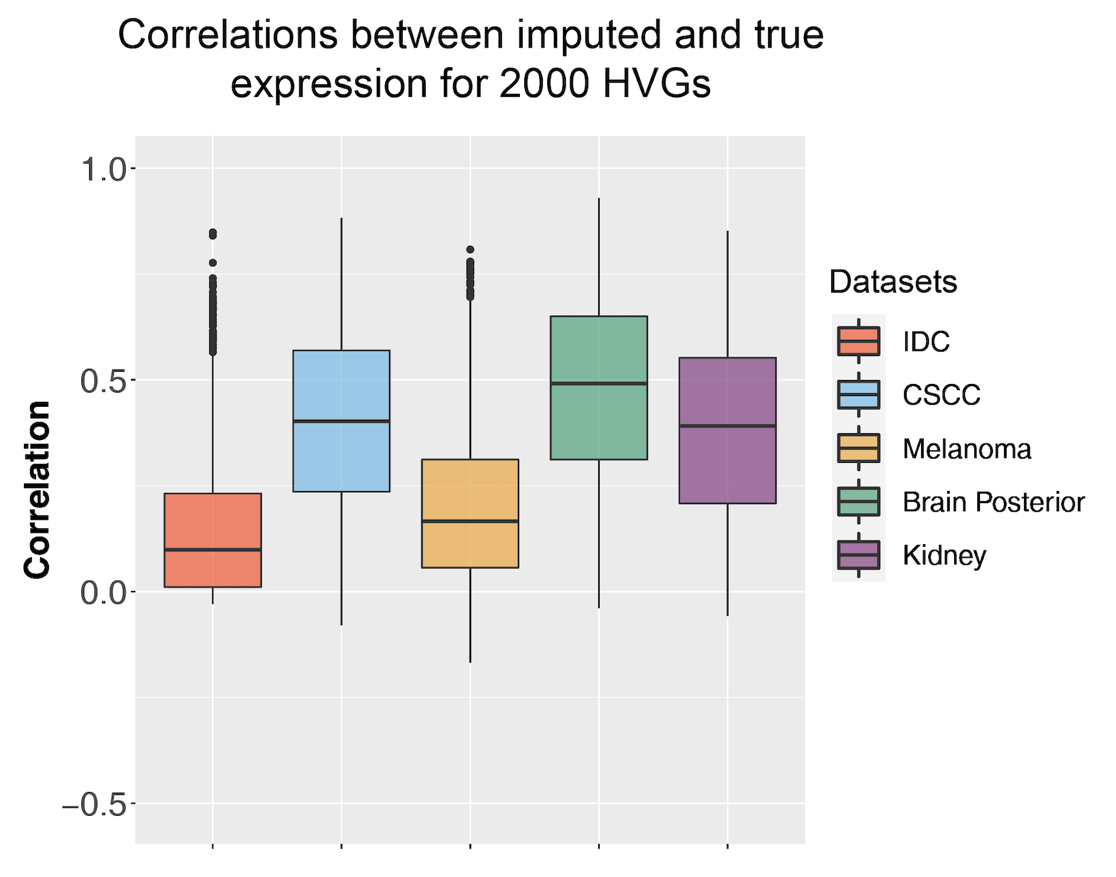

**Supplementary Fig. 2.** Enhanced gene expression by TESLA can better retain the original expression pattern at the spot level than BayesSpace. We randomly selected 10 genes in which BayesSpace’s correlation with the observed spot-level gene expression was less than 0.5.

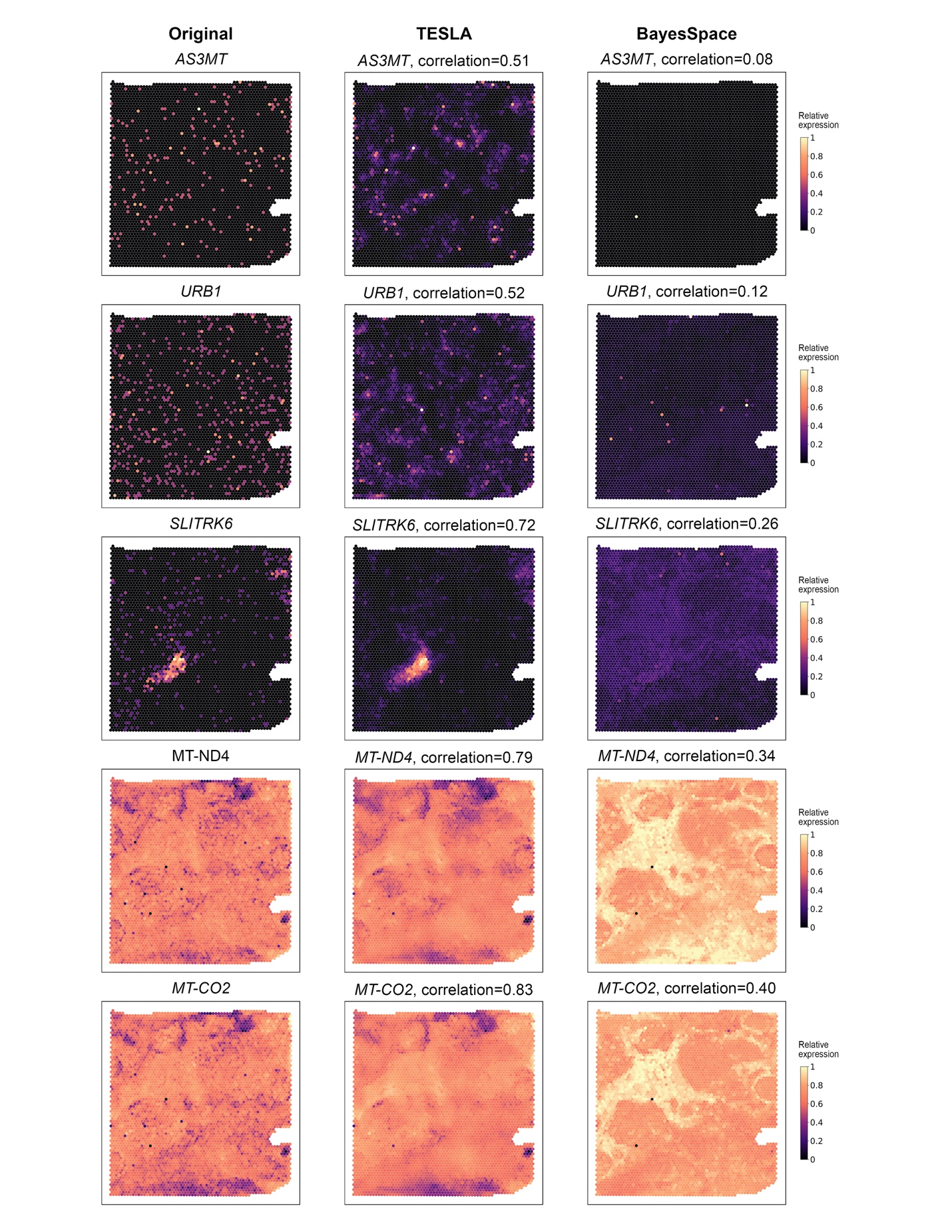

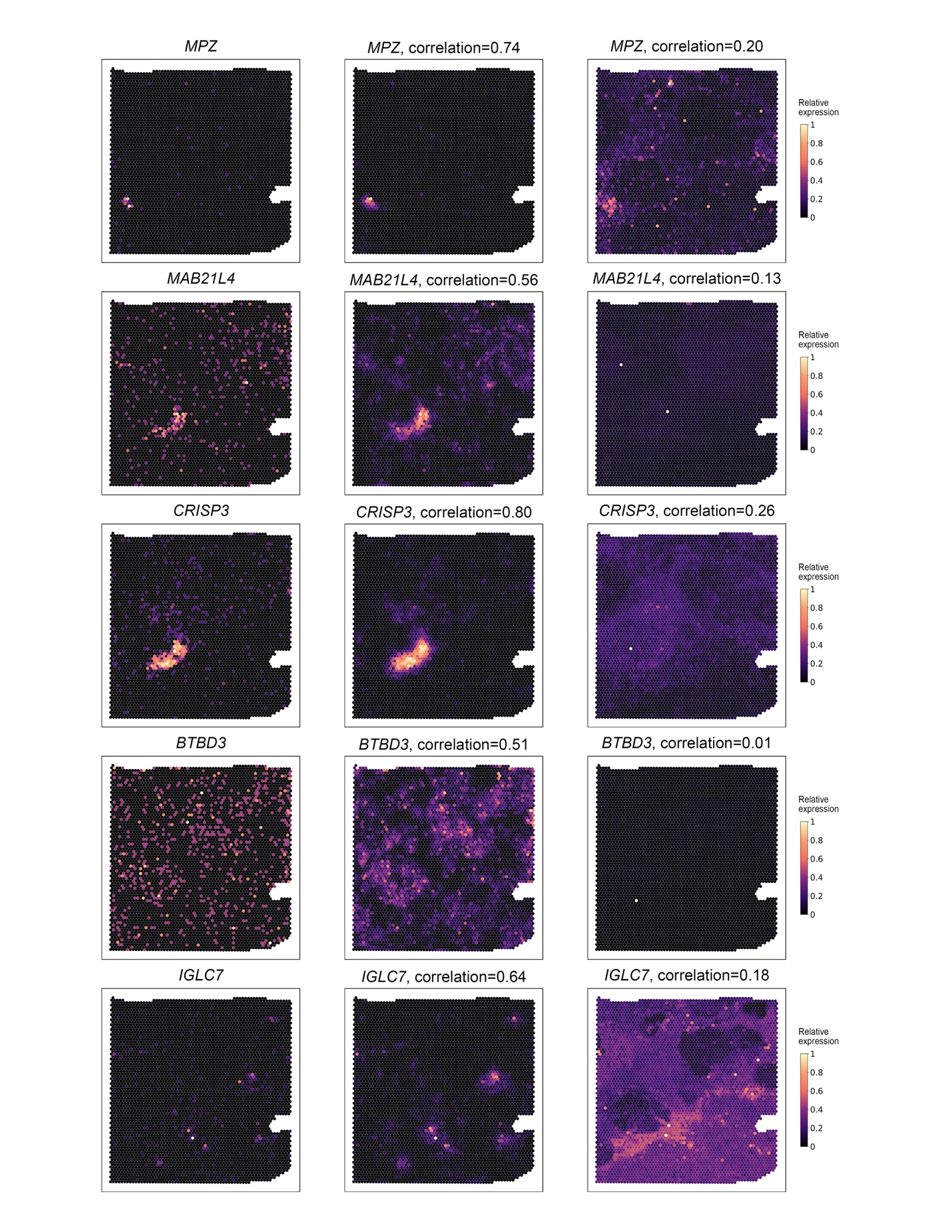

**Supplementary Fig. 3.** Comparison of enhanced gene expression accuracy by root mean squared error (RMSE) for the IDC dataset. The RMSEs were calculated for 1) TESLA enhanced *CD3E* gene expression vs CD3ε protein expression; 2) BayesSpace enhanced *CD3E* gene expression vs CD3ε protein expression; 3) TESLA enhanced *CD3E* gene expression vs CD3ε protein expression, within regions that overlap with spots; 4) BayesSpace enhanced *CD3E* gene expression vs CD3ε protein expression, within regions that overlaps with spots; and 5) the spot level RMSE for *CD3E* raw gene expression from Visium vs CD3ε protein expression as a baseline.

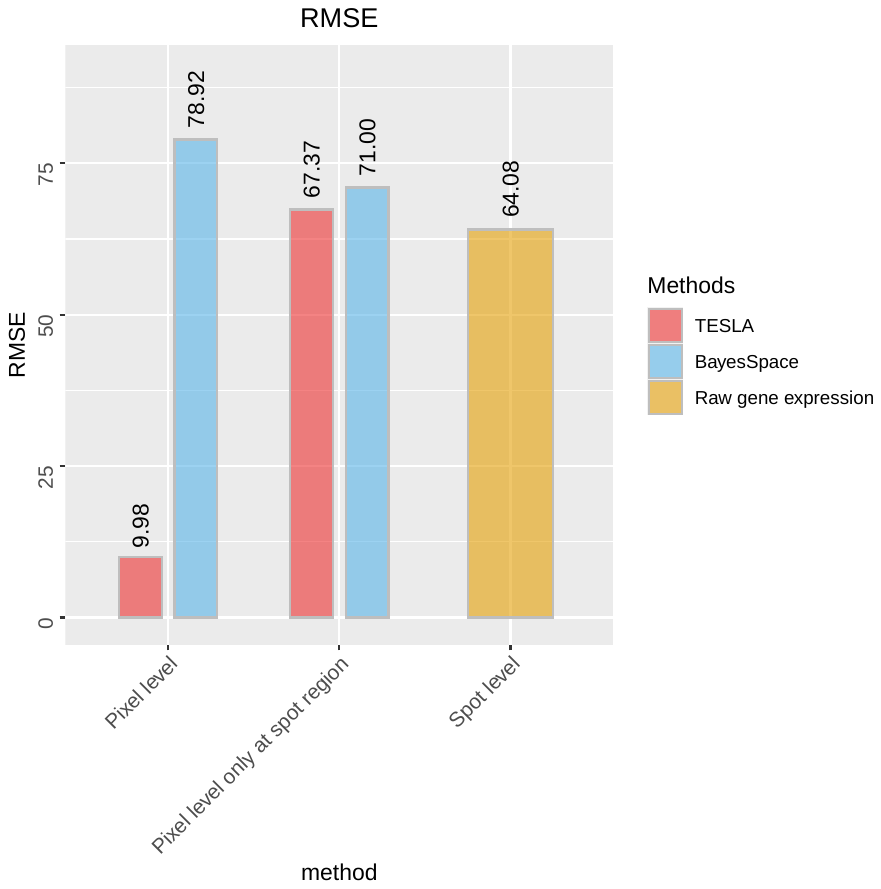

**Supplementary Fig. 4**. Tumor marker gene images for the cutaneous squamous cell carcinoma tissue section generated by TESLA.

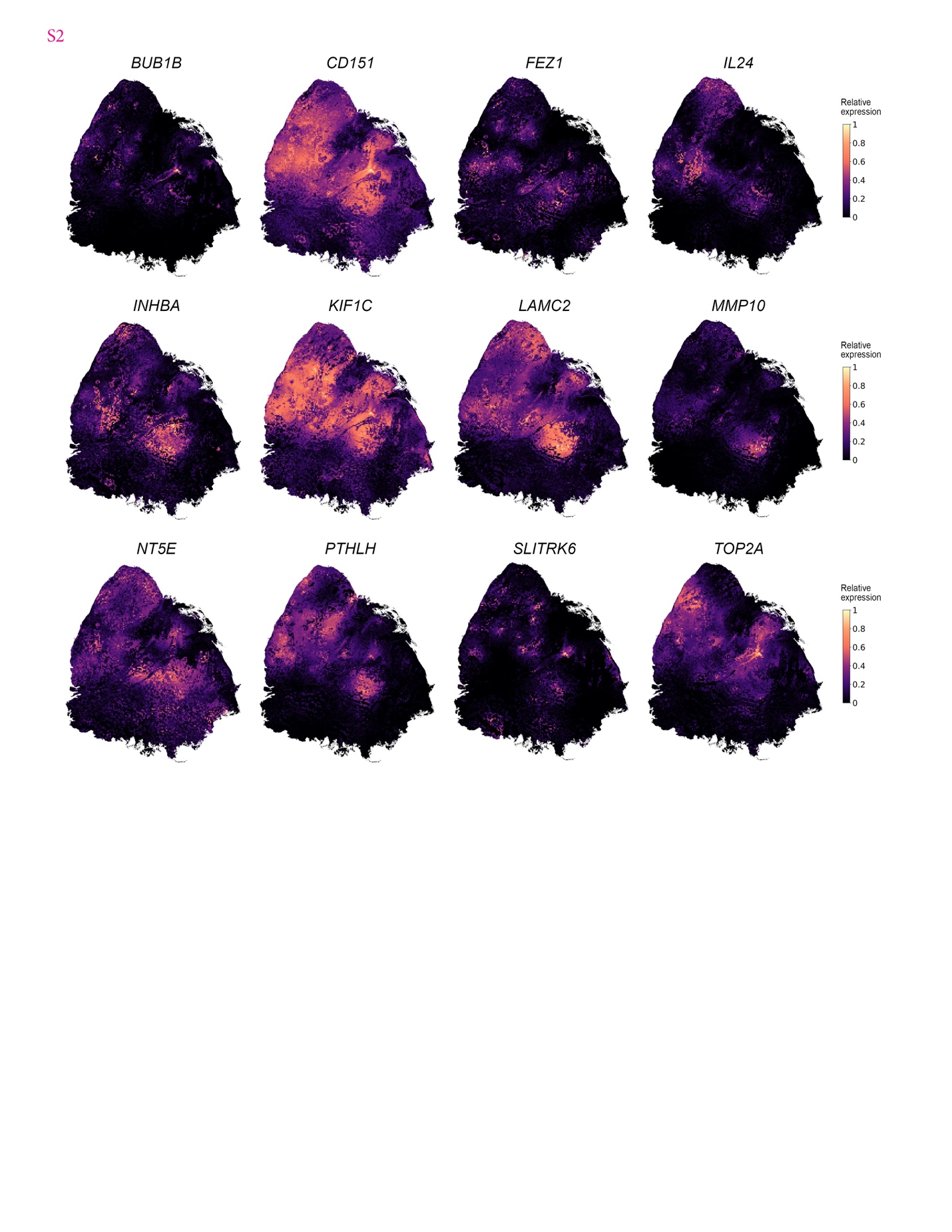

**Supplementary Fig. 5.** Total UMI counts for the cutaneous squamous cell carcinoma tissue section.

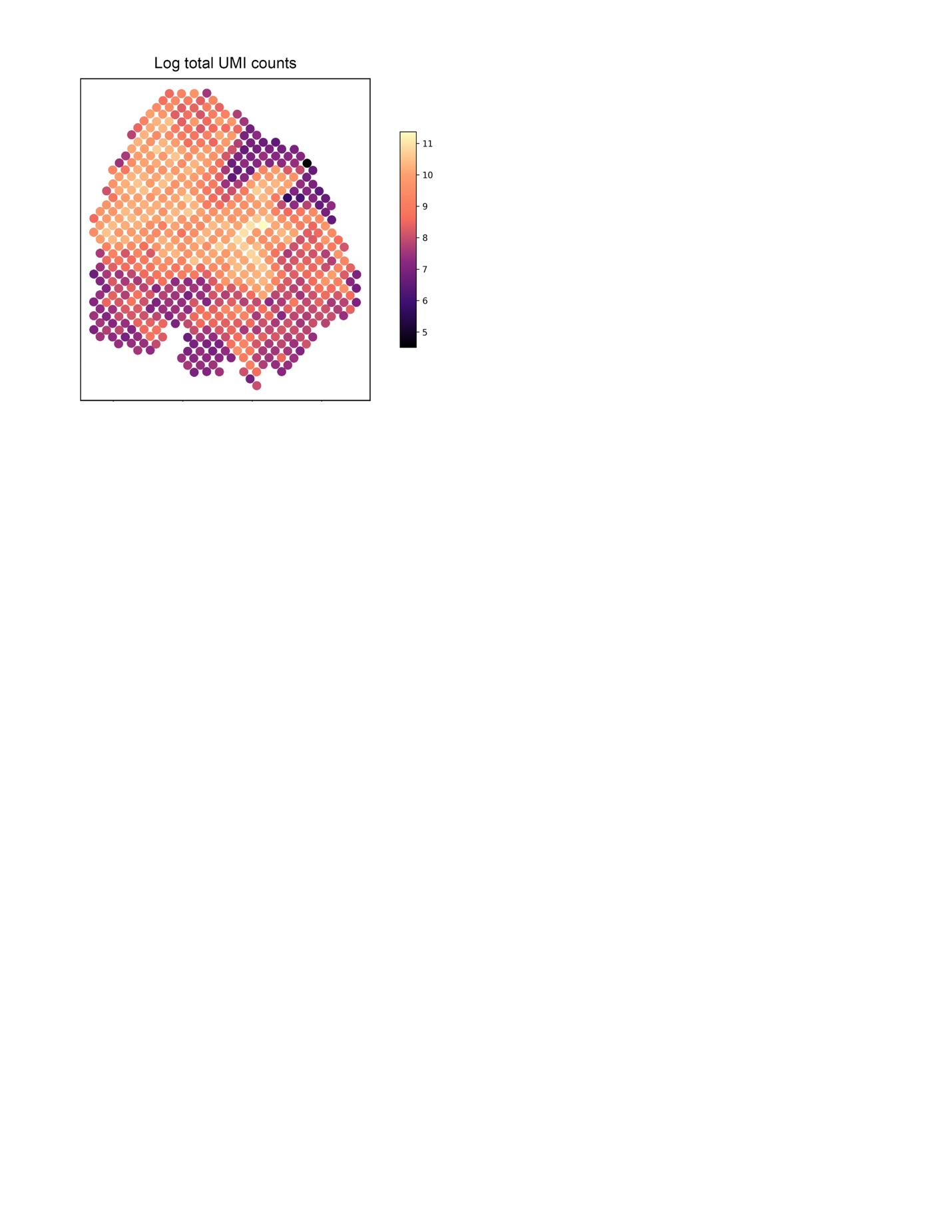

**Supplementary Fig. 6.** Examples of TESLA identified genes that are highly enriched in the tumor core or edge in the human cutaneous squamous cell carcinoma dataset of the skin.

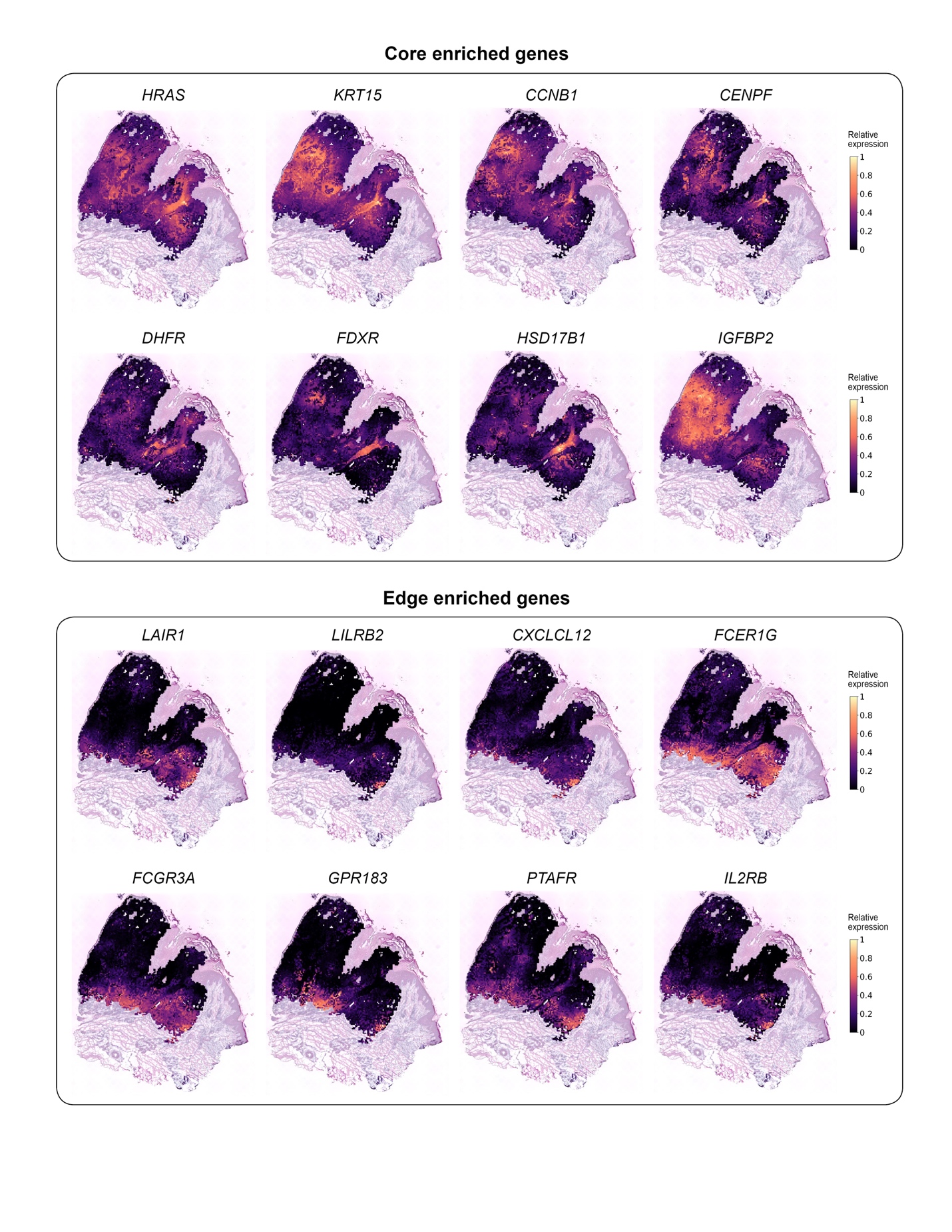

**Supplementary Fig. 7.** Marker gene images for clinical melanoma diagnosis for the cutaneous malignant melanoma tissue section.

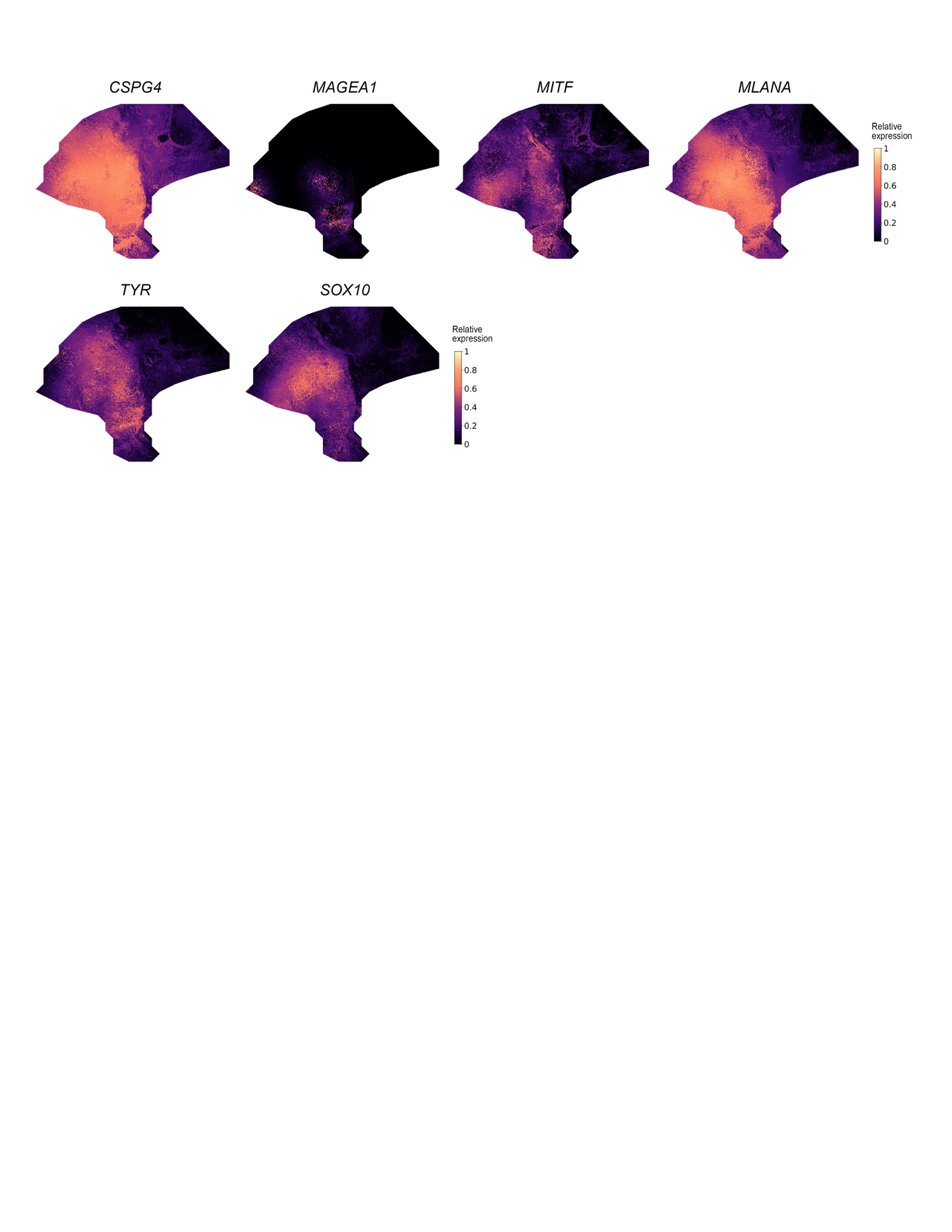

**Supplementary Fig. 8.** Examples of TESLA identified genes that are highly enriched in the tumor core or edge in the human cutaneous malignant melanoma dataset.

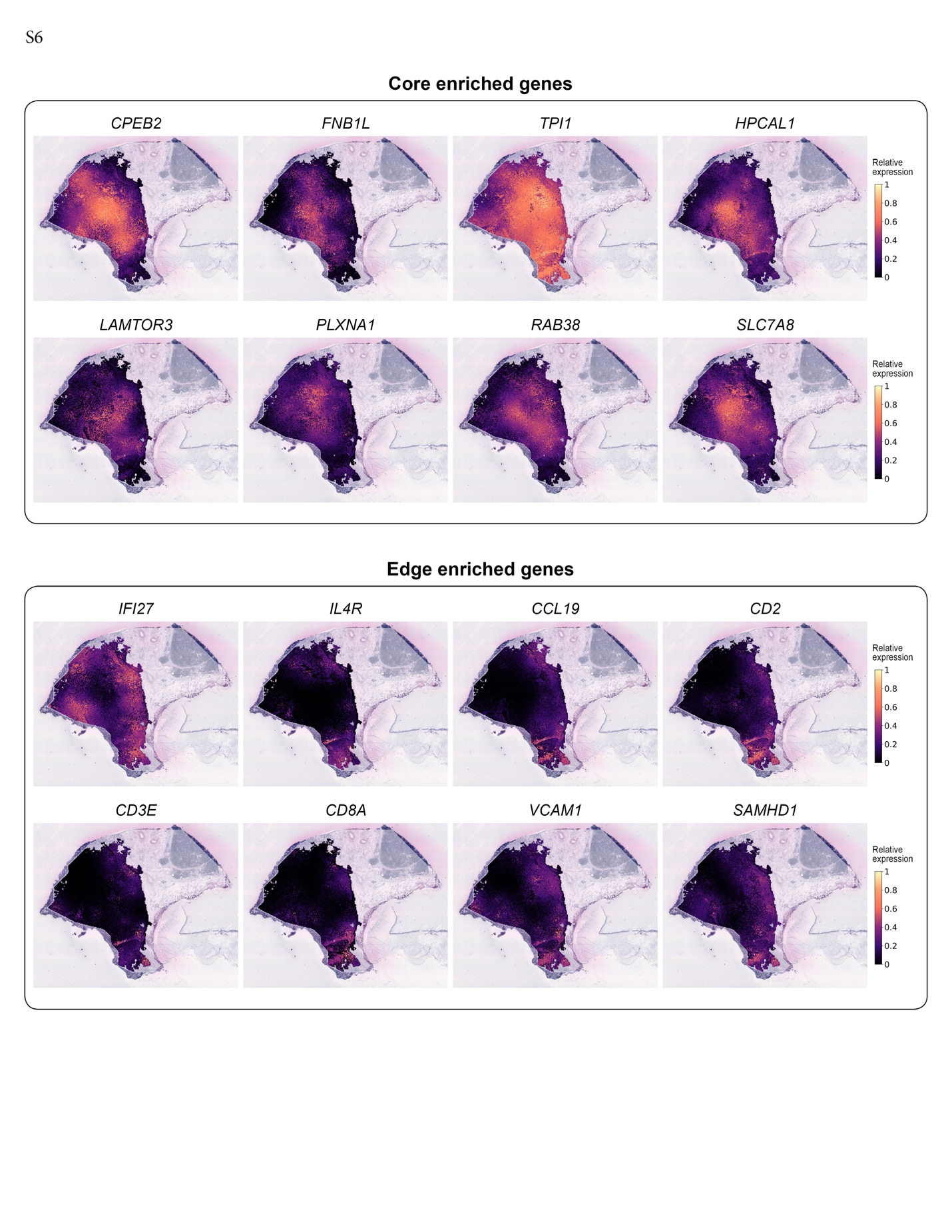

**Supplementary Fig. 9.** Cell type distributions in tumor edge and core based on deconvolution results obtained from RCTD. CAF stands for cancer associated fibroblast.

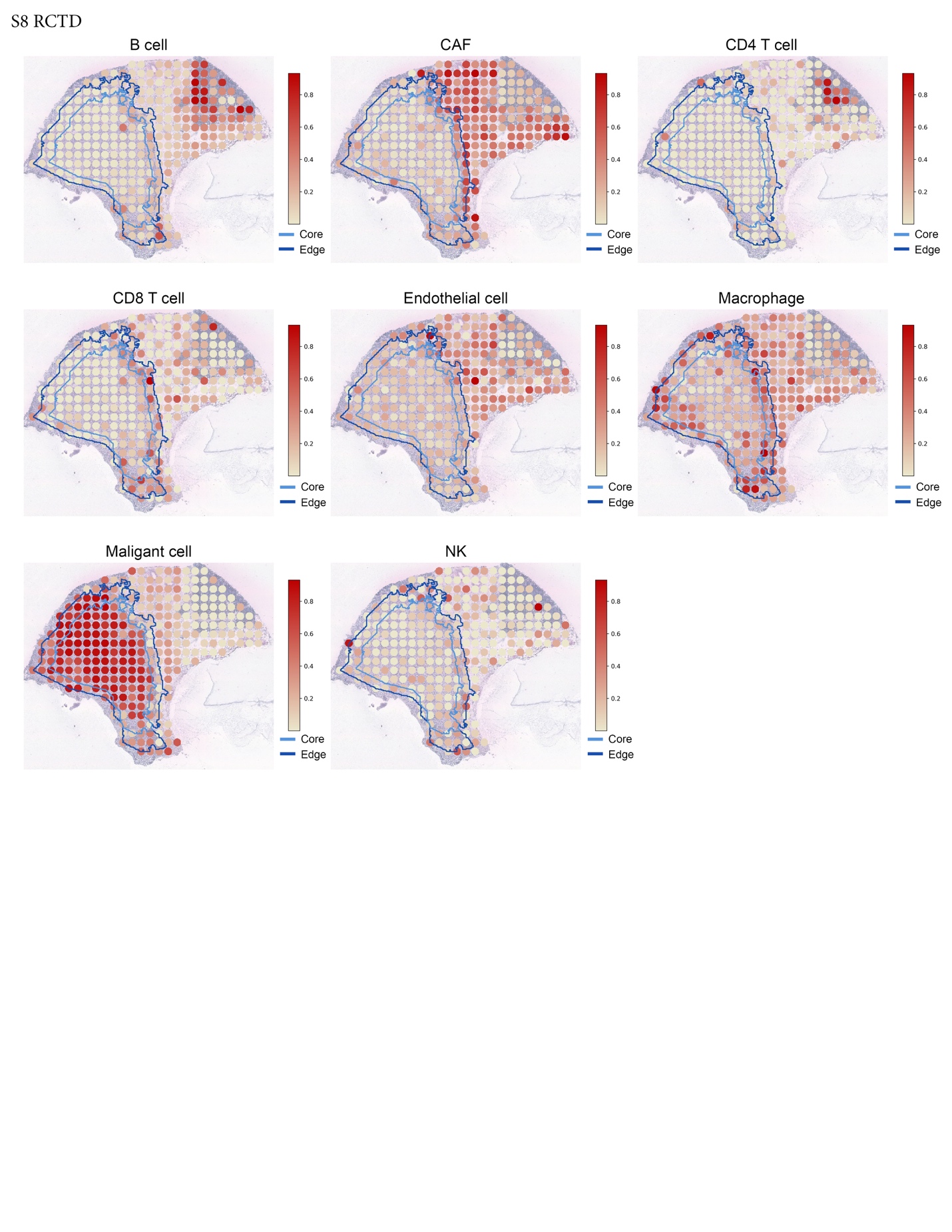

**Supplementary Fig. 10.** Marker gene images for B cells, CD4+ T cells, dendritic cells, and CXCL13 in the squamous cell skin carcinoma dataset.

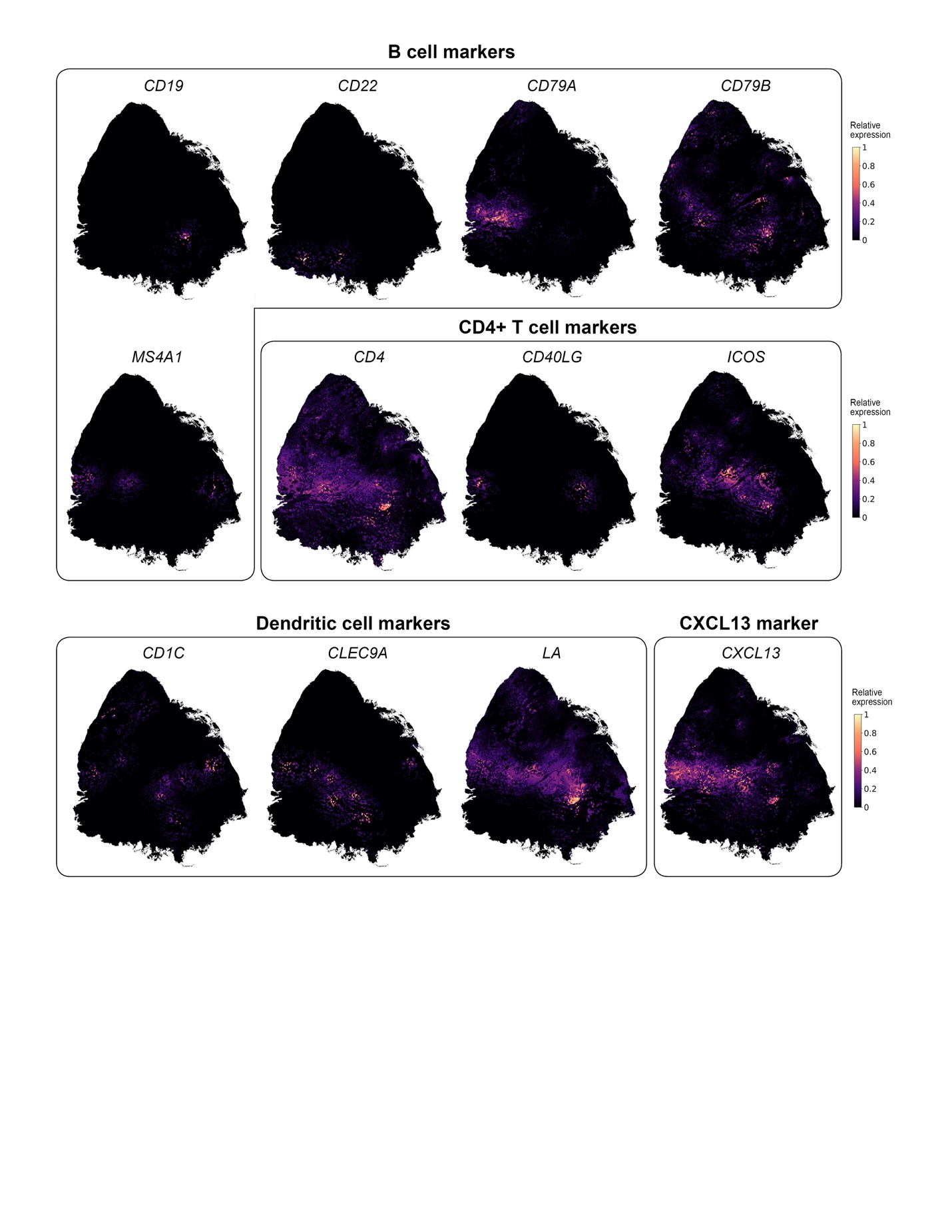

**Supplementary Fig. 11.** Marker gene images for B cells, CD4+ T cells, dendritic cells, CD8+ T cells, and CXCL13 in the human cutaneous malignant melanoma dataset.

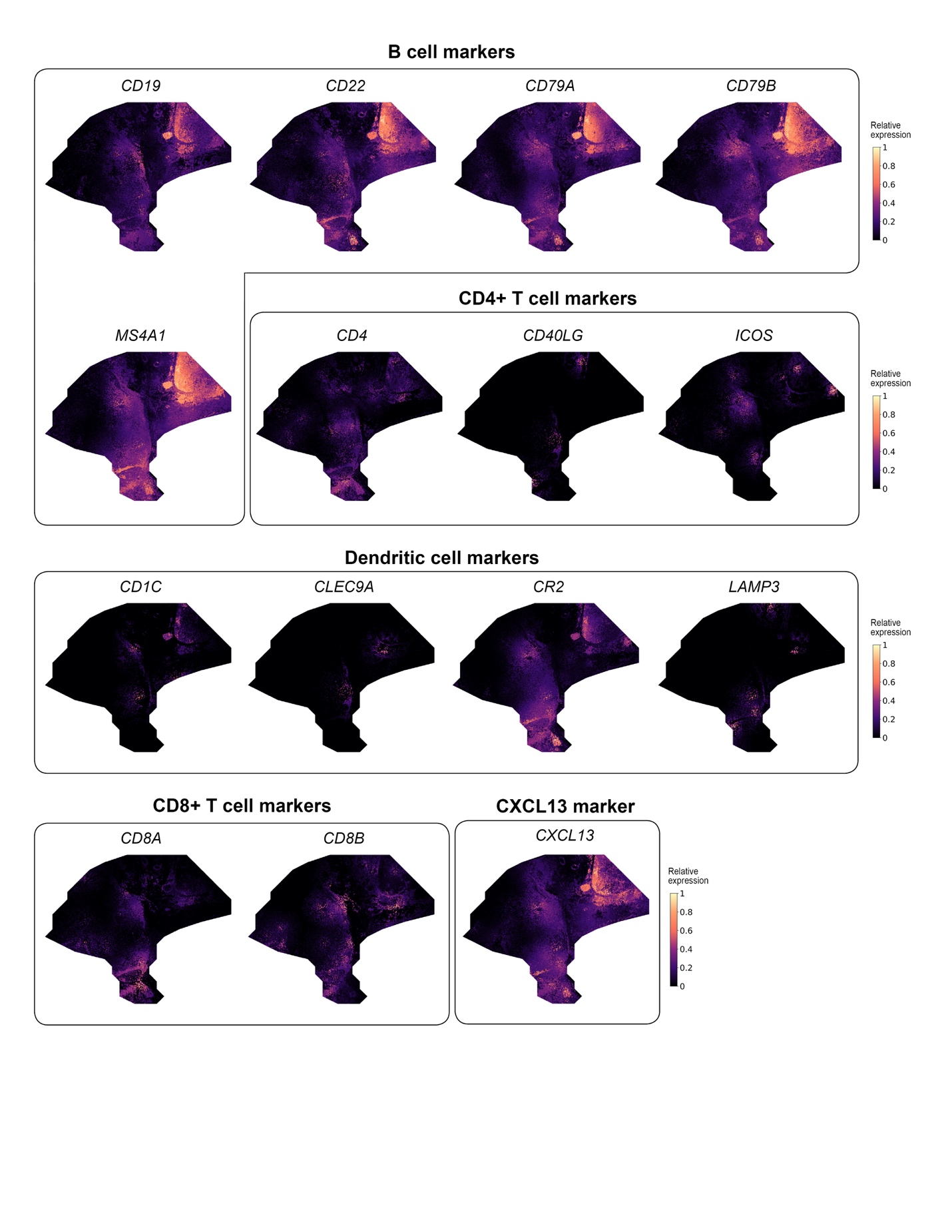

**Supplementary Fig. 12.** Cell type deconvolution results for the CSCC data using RCTD.

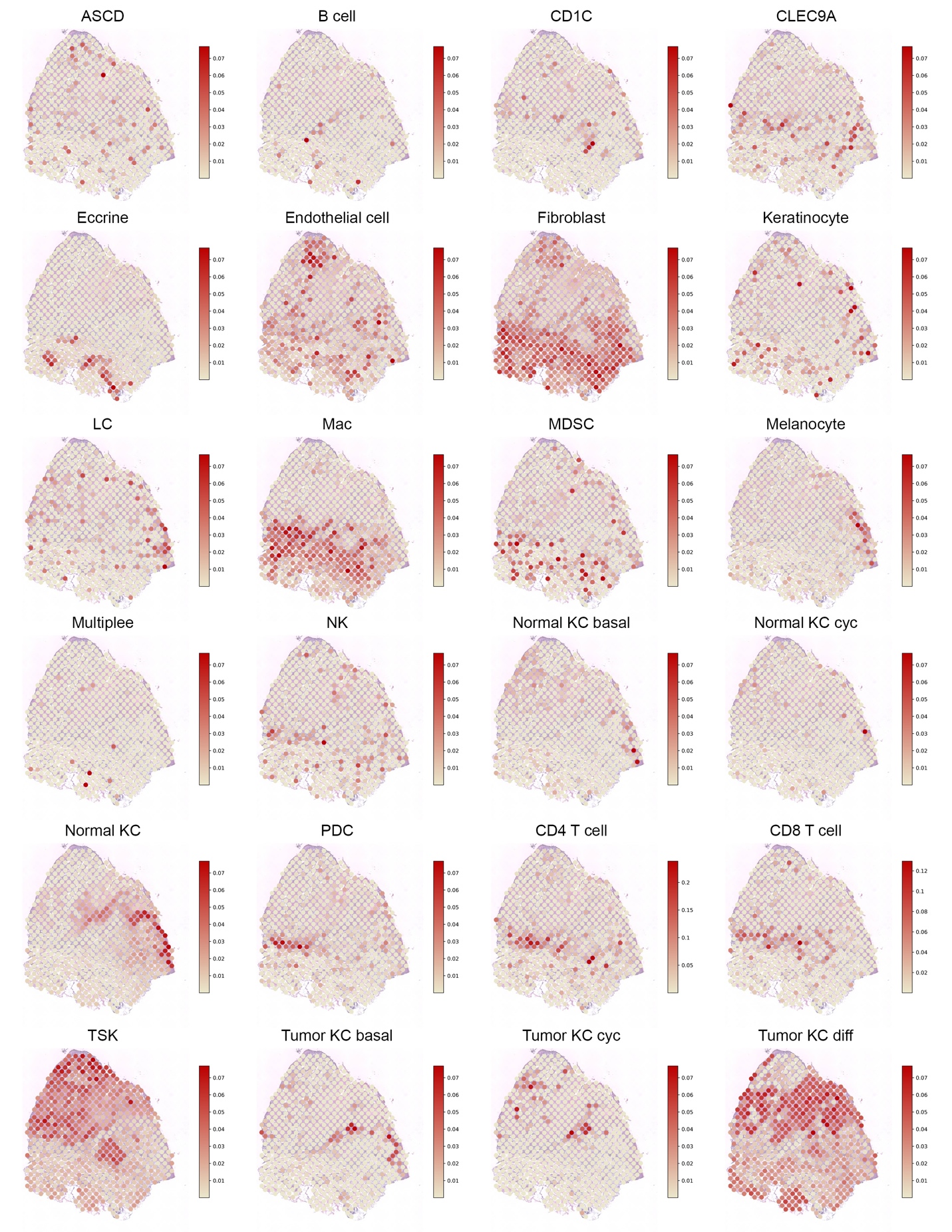

**Supplementary Fig. 13.** Cell type deconvolution results for the human cutaneous malignant melanoma data using RCTD.

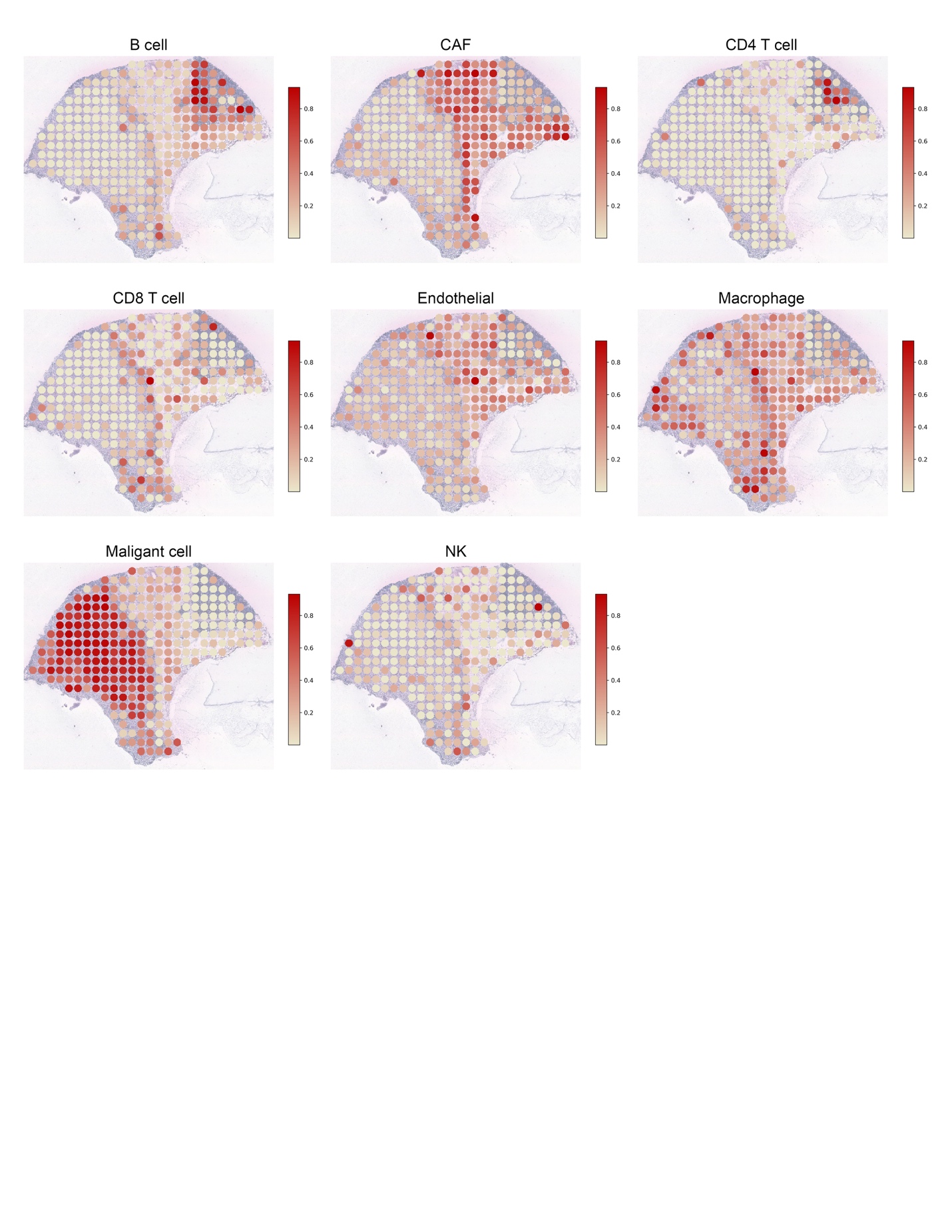

**Supplementary Fig. 14.** Distribution of follicular helper T cells in the CSCC and Melanoma data from TESLA, using markers *CD3E, CD3D, CD3G, CD4, PDCD1, CXCR5*.

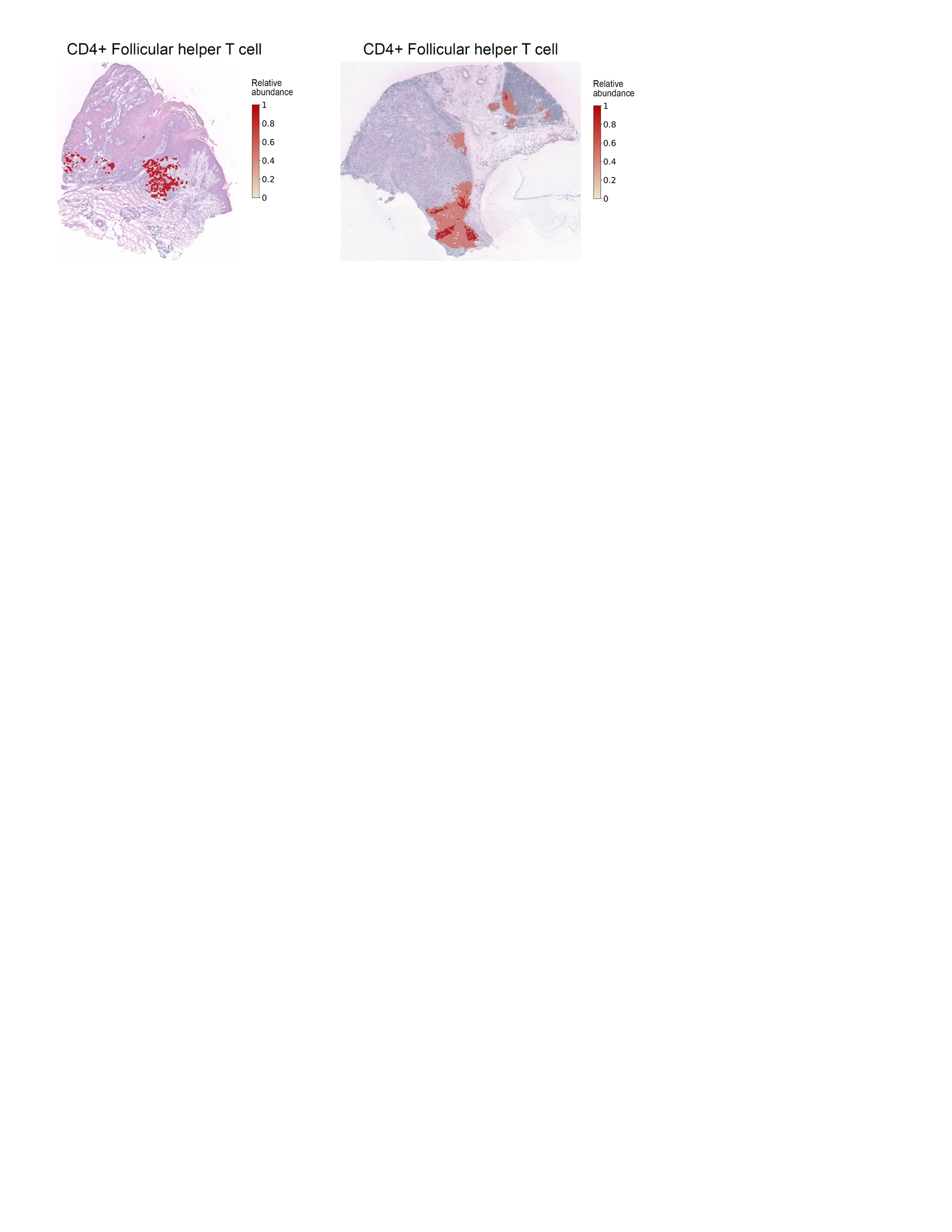

**Supplementary Fig. 15.** Tumor marker gene images in the human breast cancer datasets.

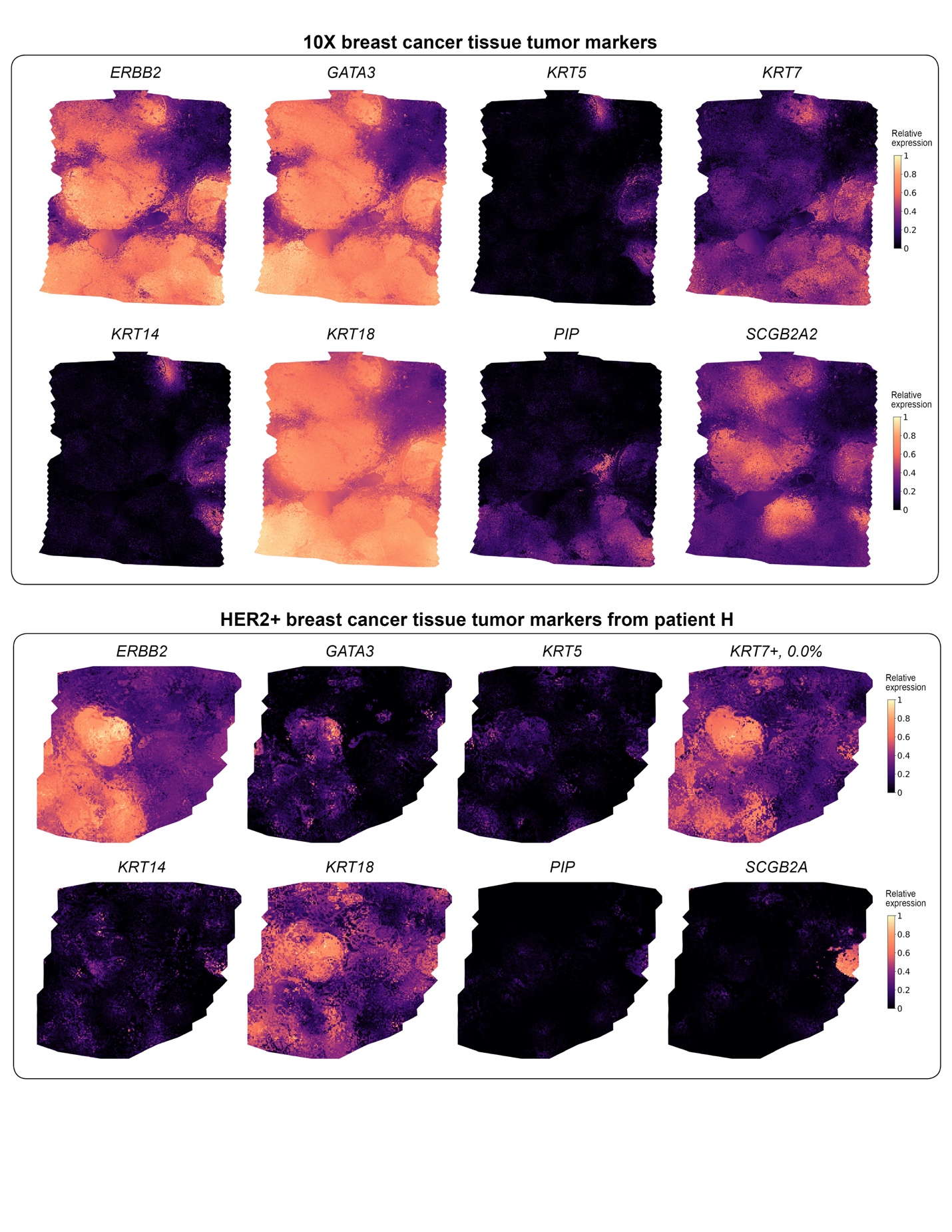

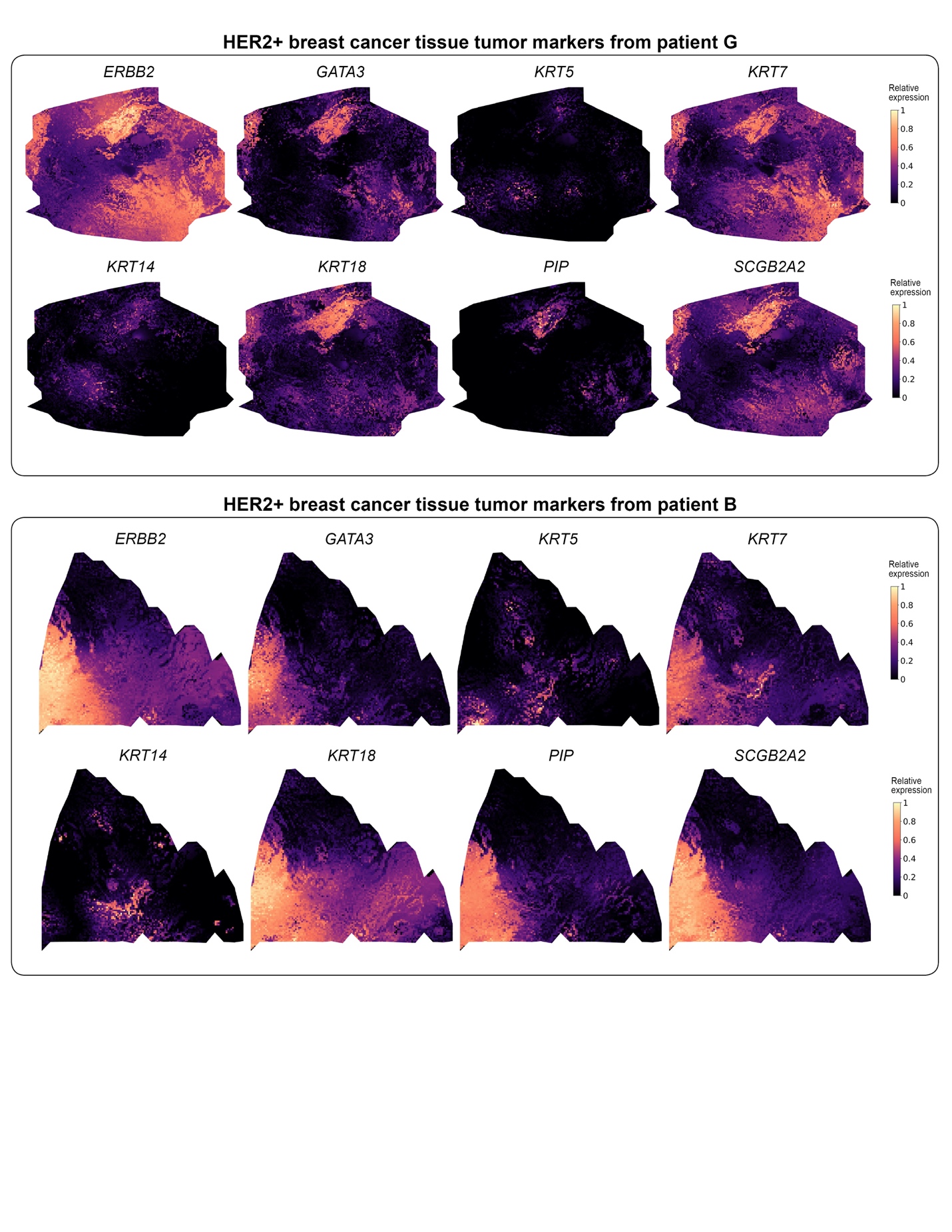

**Supplementary Fig. 16.** Super-resolution meta gene from TESLA is able to correct artifact in protein immunofluorescence staining image.

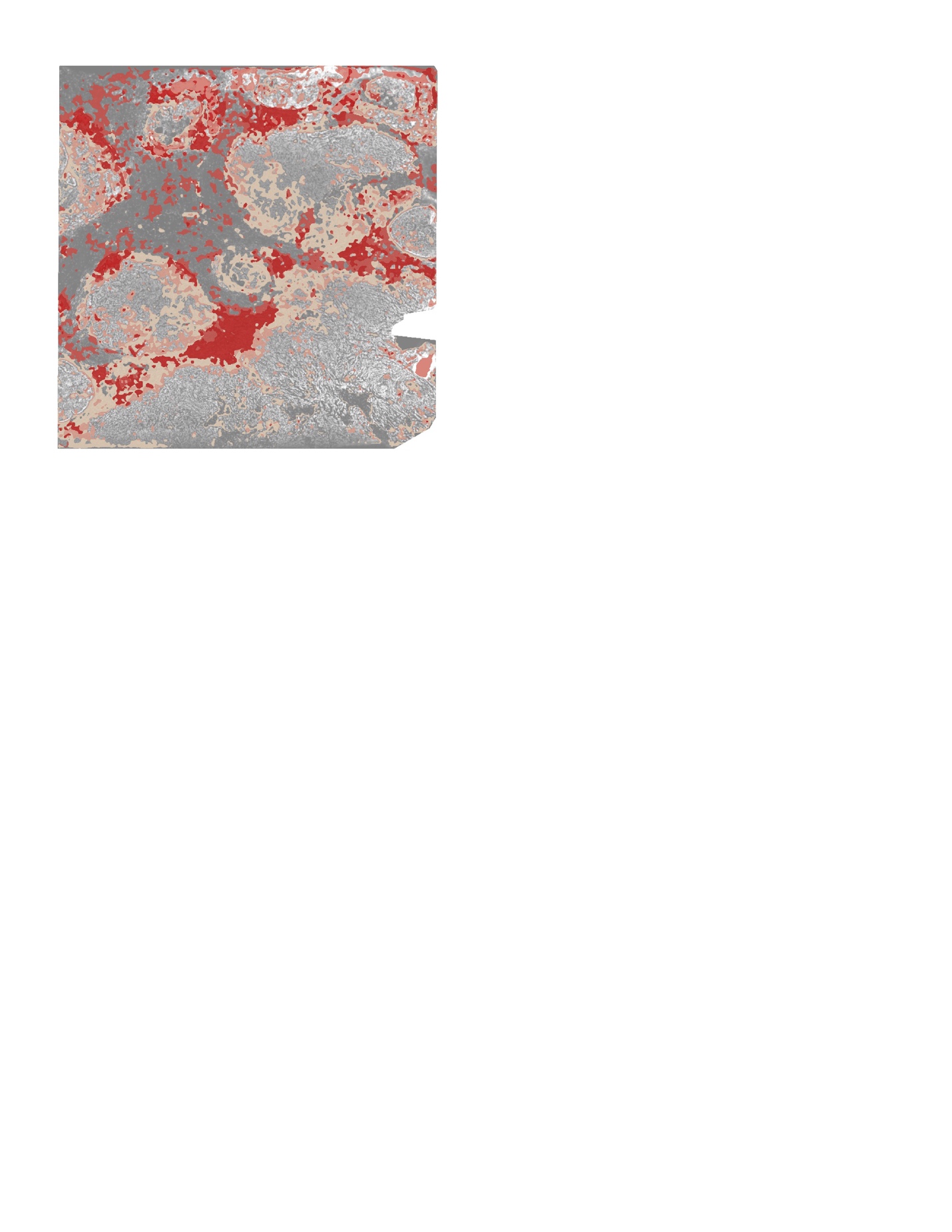

**Supplementary Note 1.** **Tissue coverage rate of different ST technologies.**

For each dataset, we separated the whole tissue area into same sized grids where each grid unit contains the same number of spots with the same pattern as shown in **Supplementary Fig. 17**. We then calculated the theoritical coverage rate using the area of spots inside the grid unit divided by the area of the grid. For 10x Visium, there are two spot layouts (10x Visium type1: invasive ductal carcinoma data; 10x Visium type2: squamous cell skin cancer cacinoma data), whose theoritical coverage rates are 25.6% and 45.8%, respectively, while for Spatial Transcriptomics (melanoma data; HER2+ breast cancer data), it has a theoritical coverage rate of 34.9%.

Additionally, we calculated the exact coverage rate for each analyzed dataset. We first detected the whole tissue region and calculated its area. Next, we derived the covered tissue area by multiplying the number of measured spots and the unit spot area. The coverage rate is computed as the ratio of covered tissue area to the whole tissue area. The invasive ductal carcinoma data from 10x Visium with pattern 1 has a coverage rate of 27.2% while the squamous cell skin cancer carcinoma data from 10x Visium with pattern 2 has a coverage rate of 48.5%. The melanoma and HER2+ breast cancer data from Spatial Transcriptomics have coverage rates of 21.4%, 20.4% (patient B), 20.3% (patient G), and 20.4% (patient H), respectively.

**Supplementary Fig. 17.** Spot layout in different ST technologies.

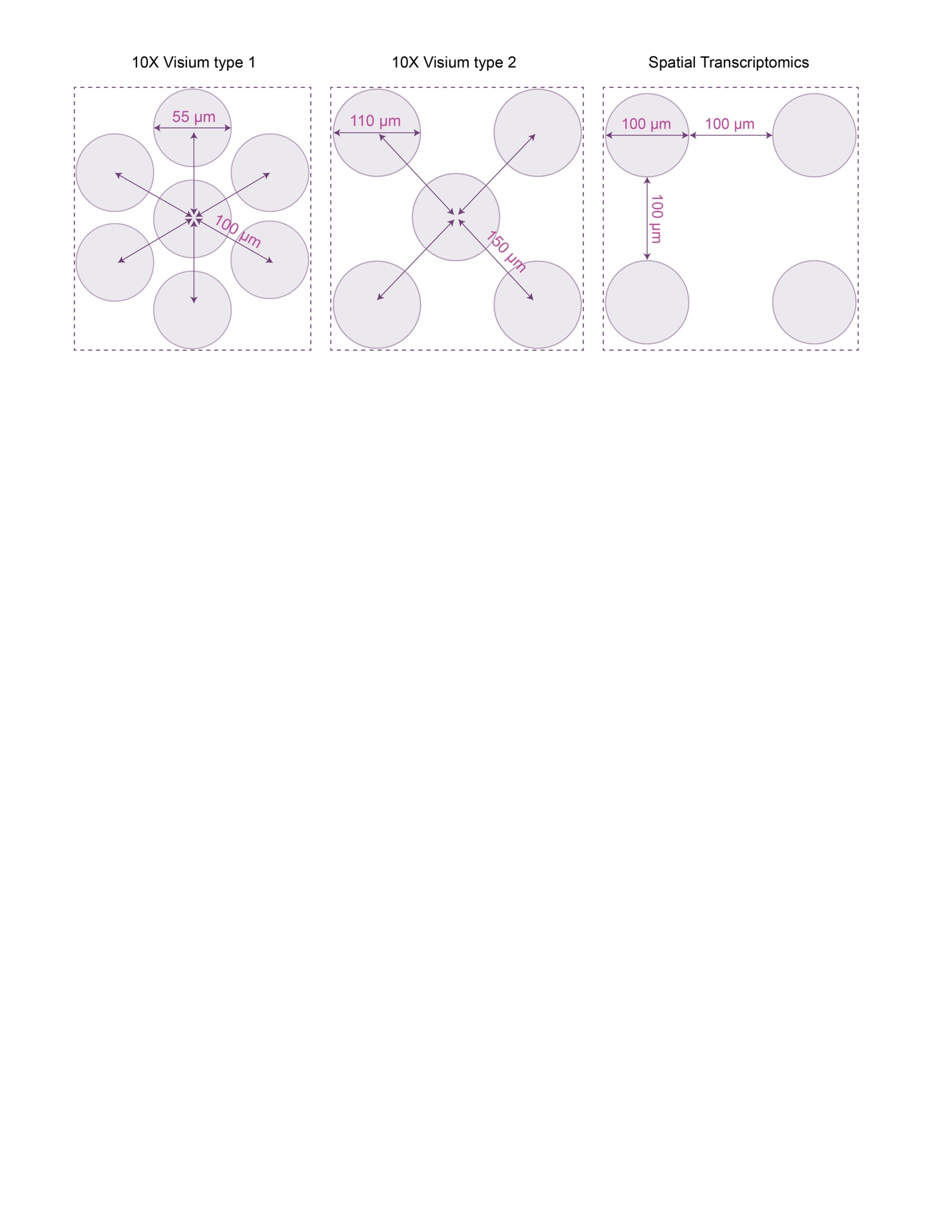

**Supplementary Note 2. The cellular and molecular spatial structure of tumor cannot be revealed with original spot-level data.**

To showcase the strength and necessity of TESLA’s super-resolution annotation, we analyzed the CSCC and melanoma datasets using their original spot-level data and compared with results obtained from TESLA. We first performed spatial clustering analysis using SpaGCN[10], a spatial clustering method that we previously developed for spatial domain detection in spatial transcriptomics. SpaGCN operates at the spot-level. We performed spatial clustering analysis with different resolution parameter values, leading to different number of clusters, i.e., spatial domains. As shown in the **Supplementary Figs. 18** and **19**, using spot-level gene expression data as input, clustering analysis cannot identify spatial domains that capture the tumor edge and core structure for both datasets, regardless how many clusters were specified in the clustering analysis.

Next, we show that the edge and core enriched genes can only be detected at the super-resolution level. To demonstrate this point, we first assigned the identity of each spot based on the tumor edge and core separation obtained from TESLA for both CSCC and cutaneous malignant melanoma data (**Supplementary Fig. 20 a,b**). Then, we performed core vs edge differential expression (DE) analysis at the spot level using the same filtering criteria as we did for the super-resolution gene expression data.

For the CSCC, we detected 3,665 genes enriched in tumor core and 106 genes enriched in tumor edge when using the super-resolution gene expression data as input, but when using the spot-level gene expression data as input, we only detected 1,023 enriched genes for the tumor core (765 genes overlap with super-resolution detection) and 0 enriched genes for the tumor edge. Similarly, for the melanoma dataset, we detected 3,510 genes enriched in tumor core and 155 genes enriched in tumor edge when using super-resolution gene expression data as input for DE analysis. But when using the spot-level gene expression as input, we only detected 1,632 genes enriched for the core (1509 genes overlap with super-resolution detection) and 1 gene (“BGN”) enriched for the edge (1 gene overlaps with super-resolution detection).

The above results show that the super-resolution gene expression data are needed to identify tumor core and tumor edge enriched genes, especially for the tumor edge. We think the failure of detecting tumor edge enriched genes at the spot level is due to two reasons. First, the number of observations is much smaller when considering spot as the analysis unit, and the reduced sample size in DE analysis will lead to less power. Second, the spot-level data do not have single-cell resolution. Indeed, the diameter size of each spot is 100um in Spatial Transcriptomics, which is much larger than a single cell. Since each spot may contain many cells, the mixture of cells from different cell types will dilute the differential expression signal, especially when the immune cells are rare. Therefore, we believe that until the sequencing-based spatial transcriptomics technologies reach to single-cell resolution, gene expression resolution enhancement will be needed when the goal is to detect gene expression changes that occur only in a small region of the tissue.

**Supplementary Fig. 18.** Spatial domains detected using SpaGCN with different numbers of domains for the CSCC data.

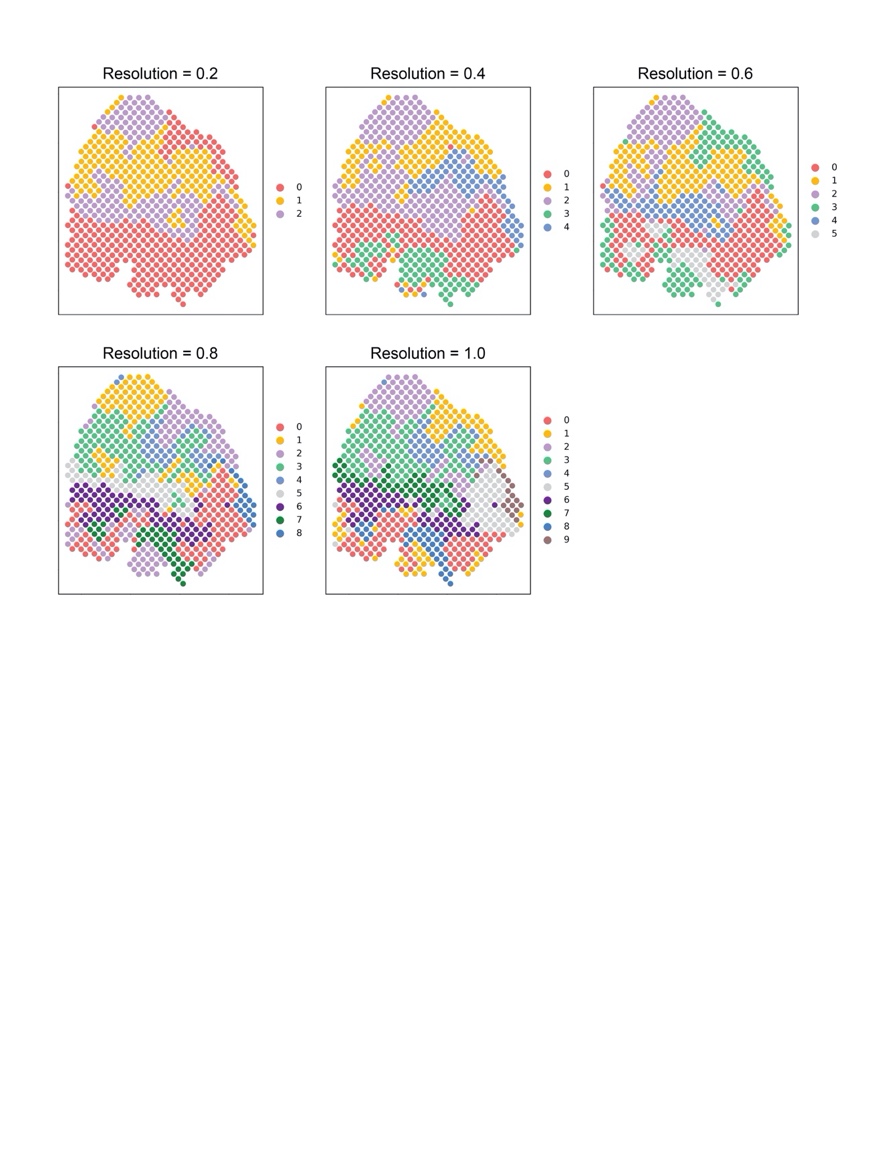

**Supplementary Fig. 19.** Spatial domains detected using SpaGCN with different numbers of domains for the melanoma data.

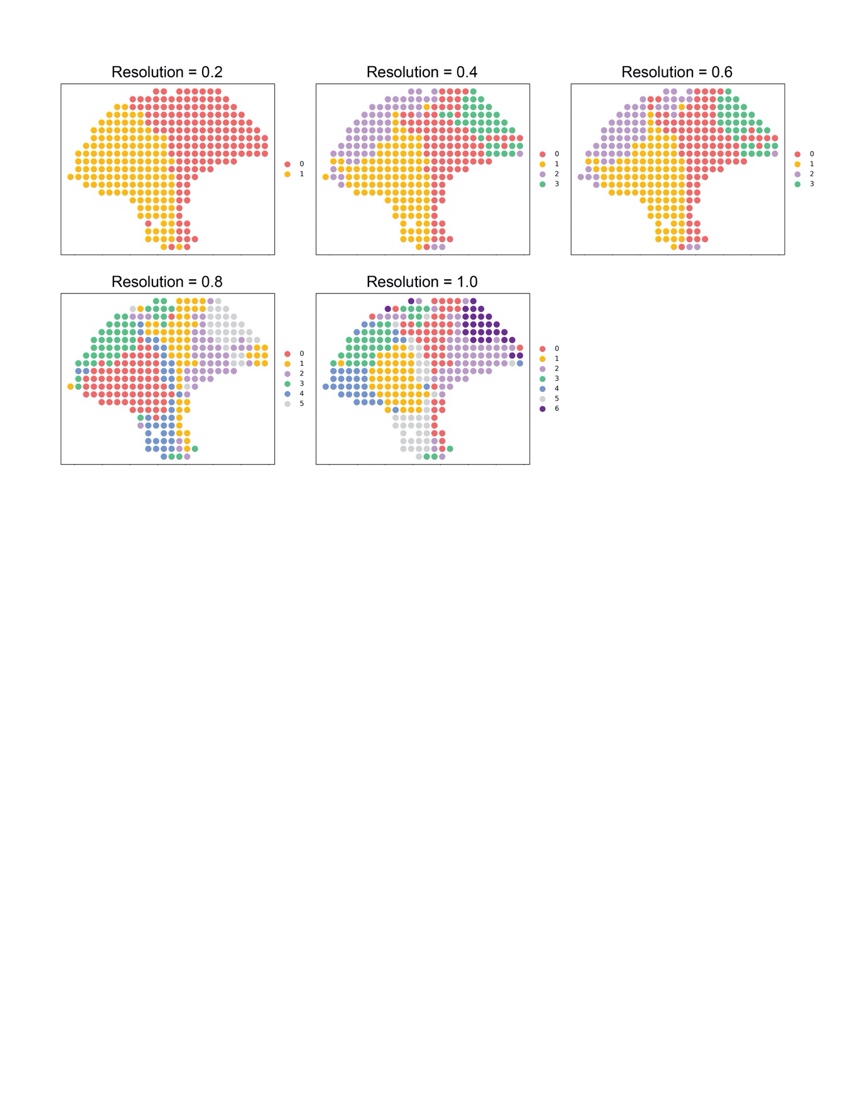

**Supplementary Fig. 20.** Spatial domains detected using SpaGCN with different numbers of domains for the melanoma data.

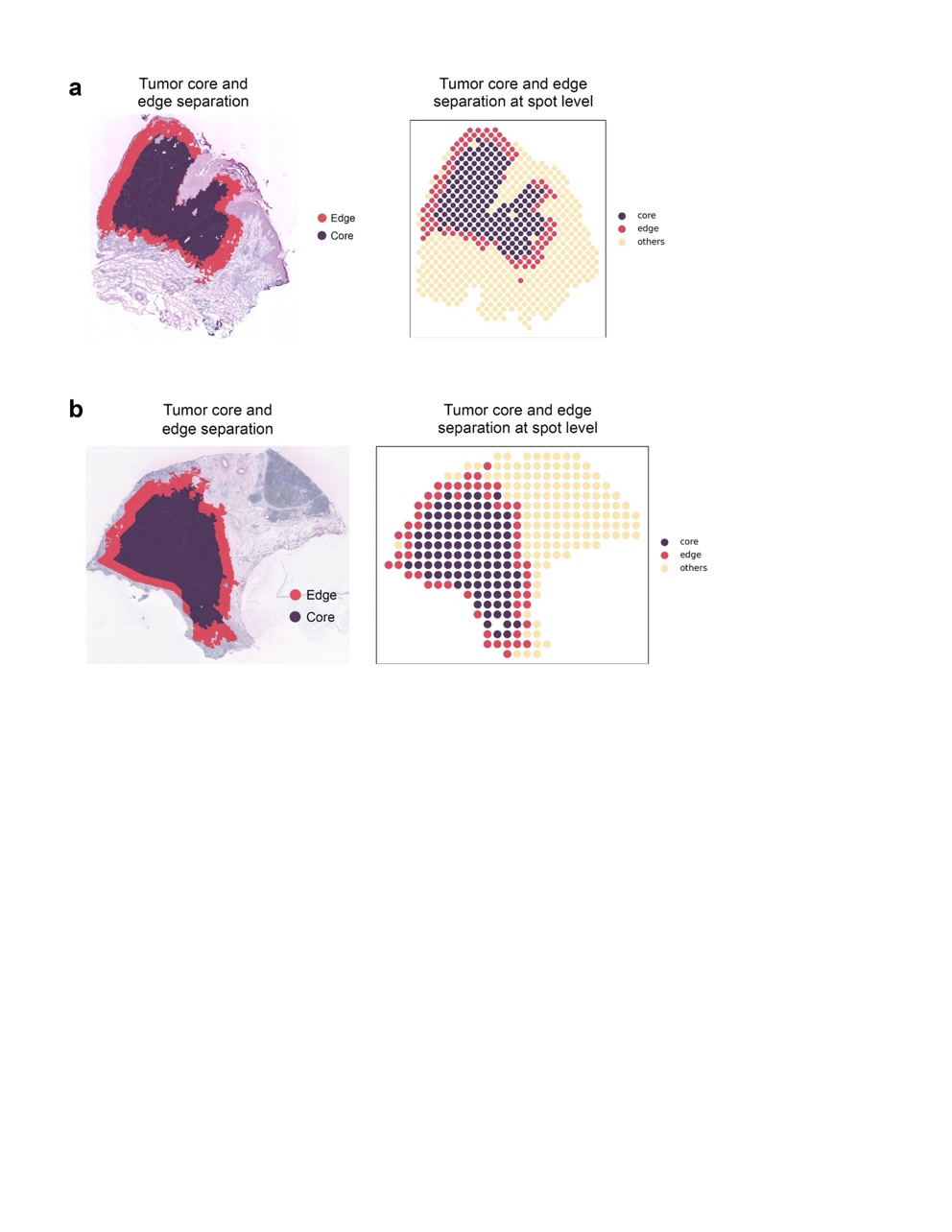

**Supplementary Note 3. Benchmark study based on Stereo-seq data.**

To establish that TESLA can recover the original high-resolution gene expression, we analyzed a Stereo-seq dataset from mouse olfactory bulb [12], which has single-cell resolution. This dataset allows us to construct a benchmark dataset where the ground truth high-resolution gene expression is known. Since the number of spots in this dataset is extremely large and the brain is symmetric, for illustration purpose, we selected the top left quarter of the brain for analysis.

We first performed clustering analysis on this dataset to identify spatial domains using SpaGCN [10]. As shown in **Supplementary Fig. 21A**, we identified 4 spatial domains. According to the annotation provided in the original study, domain 0 is the GCL region and domain 3 is the ML region. Next, we performed DE analysis between these two regions and identified 19 genes enriched in domain 0 and 14 genes enriched in domain 3 (**Supplementary Table 6**). Since Stereo-seq has single-cell resolution, we treated these DE genes as the ground truth and examined whether TESLA is able to recover these DE genes when the gene expression resolution is artificially decreased. We picked domain 0 and domain 3 for DE analysis because domain 3 is thin and without high-resolution gene expression data, it would be difficult to detect genes that are enriched in domain 3. This provides a perfect benchmark dataset that would allow us to evaluate if TESLA can help recover the enriched genes in thin tissue layers such as domain 3.

To mimic 10x Visium, we generated a benchmark dataset with artificially decreased gene expression resolution from the Stereo-seq brain data. To do so, we first merged spots in the Stereo-seq data using a grid with size 50x50 pixels, where each patch in the grid contains 0 to 7 cells, similar to the number of cells within a spot for 10x Visium. Next, we identified patches that contain cells from domain 0 or domain 3 (**Supplementary Fig. 21B**) and used these patches to perform DE analysis between domain 0 and domain 3. For domain 0, we detected 162 enriched genes with 19 overlapping with the ground truth. For domain 3, we detected 48 enriched genes with 4 overlapping with the ground truth (**Supplementary Table 6**). This analysis suggests that when analyzing the “lower-resolution” data, the true positive rate for DE analysis is low but the false positive rate is high for domain 3.

Next, we enhanced the gene expression resolution for the above benchmark data using TESLA with superpixel size 25x25 pixels. We identified superpixels that contain cells from domain 0 and domain 3 (**Supplementary Fig. 21C**), and then used these superpixels to perform DE analysis. For domain 0, we detected 23 enriched genes with 19 overlapping with the ground truth. For domain 3, we detected 20 enriched genes with 12 overlapping with the ground truth (**Supplementary Table 6**). Compared with the results obtained from the benchmark data without gene expression resolution enhancement, we substantially improved true positive rate and decreased false positive rate, suggesting that TESLA is able to recover biologically meaningful super-resolution gene expression.

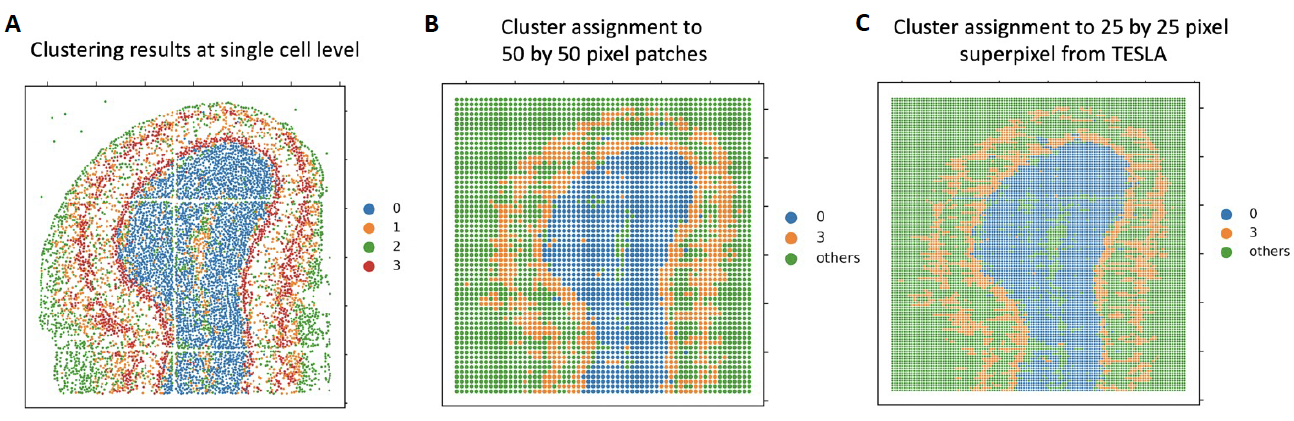

**Supplementary Fig. 21: A,** Clustering results at the single-cell level in the original Stereo-seq data (left); **B,** 50x50 pixel patches that correspond to domain 0 (blue) or domain 3 (orange); **C**, 25x25 pixel superpixels from TESLA that correspond to domain 0 (blue) or domain 3 (orange).

**Supplementary Table 6:** Number of enriched genes for domain 0 and domain 3.

|  | Domain 0 | Domain 3 |
| --- | --- | --- |
| Ground truth (Stereo-seq) | 19 enriched genes | 14 enriched genes |
| 50x50 pixel patches (mimic 10x Visium) | 162 enriched genes (19 true) | 48 enriched genes (3 true) |
| 25x25 pixel superpixels from TESLA | 23 enriched genes (19 true) | 20 enriched genes (12 true) |

Results from this benchmark dataset demonstrate that TESLA is able to detect true DE genes, especially for domain 3, which is similar to tumor edge.

**Supplementary Note 4. Computation cost of TESLA, BayesSpace on the IDC dataset.**

TESLA is computationally fast and memory efficient. To showcase the computational advantage of TESLA, we recorded its run time and memory usage for the IDC data and compared with BayesSpace. All analyses were conducted on Mac OS 10.13.6 with single Intel® Core(TM) i5-8259U CPU @2.30GHz and 16GB memory. As shown in **Supplementary Fig. 21**, TESLA completed gene expression enhancement in 19 minutes, whereas the computing time was more than 11 hours for BayesSpace. Furthermore, TESLA only required 10.0GB of memory, whereas BayesSpace required 12.9 GB of memory.

**Supplementary Fig. 21.** TESLA and BayesSpace time and memory usage comparison using the IDC dataset.

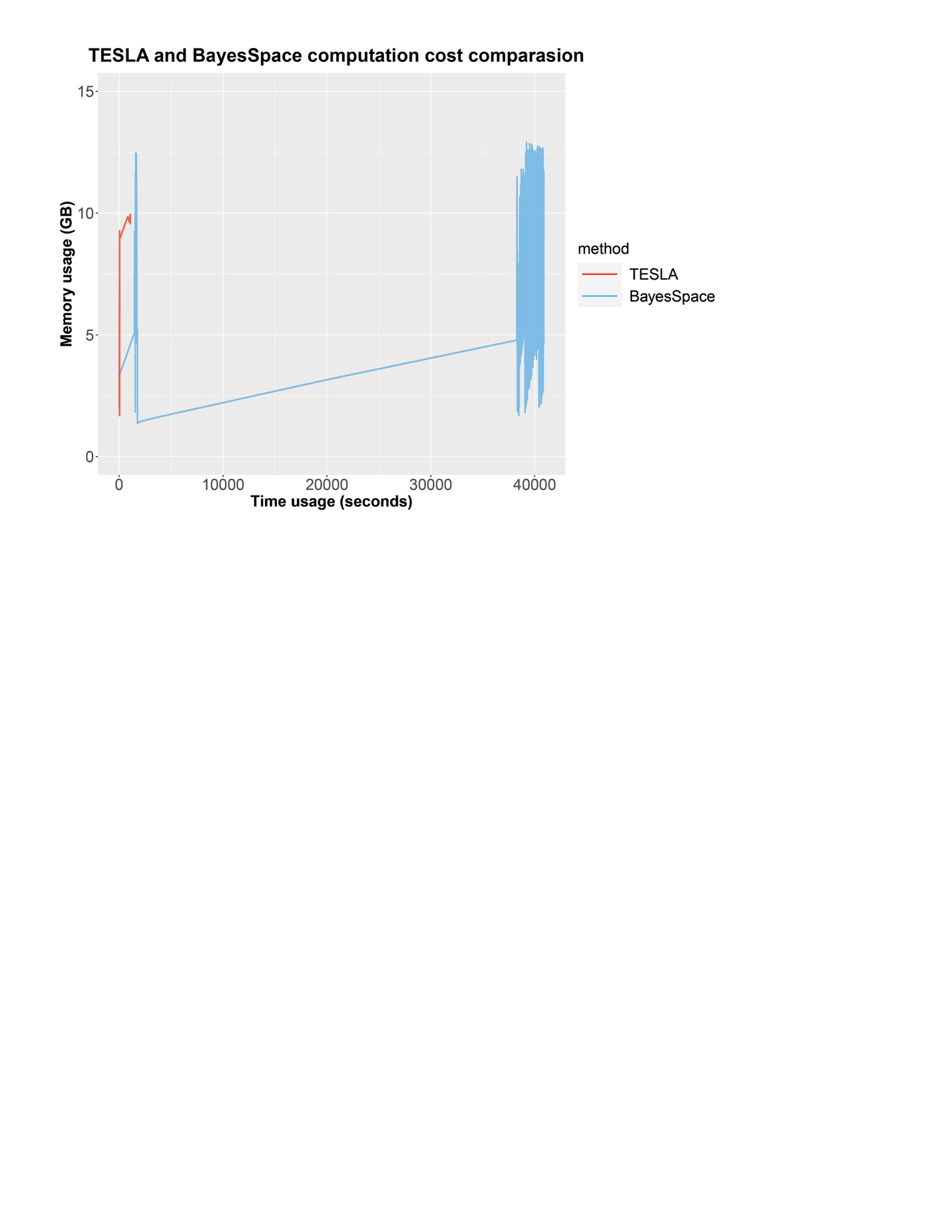
